# Reduced enteric *Bdnf* expression and TrkB-dependent regulation of *Vip* in a *Mecp2-null* mouse model of Rett syndrome-associated gastrointestinal dysmotility

**DOI:** 10.64898/2026.04.12.718037

**Authors:** Srinivas N. Puttapaka, Israel A. Admasu, Ainsleigh Scott, Gamze Sonmez, Philippa Seika, Mahalakshmi Rajkumar, Xavier Valencia, Alan Consorti, Su Min Hong, Jared Slosberg, Michela Fagiolini, Subhash Kulkarni

## Abstract

Gastrointestinal (GI) dysmotility is a prevalent and debilitating feature of Rett syndrome (RTT), yet its underlying mechanisms remain poorly understood. We investigated enteric mechanisms of dysmotility in *Mecp2-null* male mice. MeCP2 was expressed in myenteric ganglia, including enteric neurons. Reduced expression of the dominant enteric *Bdnf* transcripts, *IV* and *VI*, was evident at postnatal day 35, preceding detectable delayed transit and *Vip* downregulation at day 55. Neither TrkB.FL nor TrkB.T1 transcript expression differed significantly between genotypes at any age. In dysmotile mice, while overall neuronal density, nitrergic neuronal abundance, and *Nos1, Chat*, and *Uchl1* expression were not significantly altered, dysmotility was associated with decreased expression of *Vip* and *Cartpt* and increased expression of *Vipr1* and *Vipr2*. Interpretation alongside public single-nucleus transcriptomic data suggested selective alterations in inhibitory neuronal phenotypes or population composition. Conditional deletion of TrkB.FL in neural crest-derived cells reduced enteric *Vip* expression without reproducing the *Cartpt* or VIP receptor changes observed following MeCP2 loss. In differentiating postnatal ENS cultures from wildtype mice, pharmacological blockage of TrkB using ANA-12 reduced *Vip* and *Cartpt* expression without significantly altering *Nos1* expression. Conversely, pharmacological stimulation of TrkB using HIOC increased *Vip* expression in *Mecp2-null* cultures that otherwise showed loss of *Vip* gene expression compared to cells from wildtype mice. Together, these findings associate early enteric *Bdnf* reductions with later VIPergic abnormalities and delayed transit, while demonstrating pharmacological responsiveness of *Vip* expression to TrkB modulation. These observations support further investigation of TrkB stimulation as a potential strategy for improving enteric function in RTT.

## Introduction

Rett Syndrome (RTT) is a severe neurodevelopmental disorder, where patients undergo a developmental arrest and regression following an early period of normal post-natal development primarily caused by mutations in the X-linked gene encoding the transcriptional regulator Methyl-CpG-binding protein 2 (MeCP2) [1–3]. After the first year of life, where they undergo relatively normal development with subtle motor coordination deficiencies, RTT patients suffer from arrested neurological development leading to a regression of motor skills and loss of acquired verbal skills [3]. Apart from cognitive and motor dysfunction, RTT patients suffer from significant autonomic dysfunction. Principal amongst them are gastrointestinal (GI) dysfunctions, which manifest often as GI dysmotility disorders leading to slowed intestinal transit and chronic constipation [4]. More than 90% of RTT patients were reported to suffer from GI dysmotility disorders that severely impact nutritional status of these patients and thus could help explain the linear growth deficits, poor weight gain, and decreased bone mineralization that often characterize the clinical course of RTT patients [4]. However, despite its prevalence in RTT patients and its clinical importance, the pathobiological mechanism that causes GI dysmotility in RTT patients has remained unclear. This gap in our understanding has negatively impacted our ability to develop novel therapeutic regimens for improving GI motility in RTT patients.

GI motility is a complex, highly coordinated physiological process that is dependent on the actions of the cells of the enteric nervous system (ENS) to regulate specific motor patterns required for mixing luminal contents to aid in their digestion, and peristaltic motility required for propulsion or forward movement of luminal contents through the GI tract [5]. The ENS is the largest component of the autonomic nervous system, and GI dysmotility disorders can often be traced to ENS dysfunctions [6]. Thus, the prevalence of GI dysmotility in RTT patients suggests an underlying ENS dysfunction. Prior studies have not only documented MeCP2 expression in the ENS but also reported gastrointestinal dysmotility in an established MeCP2 knockout model of RTT (*Mecp2-null*) [7, 8]. GI dysmotility in *Mecp2-null* was associated with reduced gene expression of nitrergic neuronal marker *Nos1* in cultured ENS cells compared to those from wildtype (WT) mice and a paradoxical increase in NOS1 protein in lysates of the myenteric plexus compared to those from WT mice [7]. These observations underscore that key questions regarding whether and how ENS architecture and function are altered in this RTT model remain unanswered. Notably, these studies also observed that enteric MeCP2 expression did not alter between various fetal and early post-natal timepoints [8]. This raises another important unresolved question on whether MeCP2 loss–driven GI dysmotility emerges before, or in parallel with, the onset of developmental arrest in RTT.

In the Central Nervous System (CNS), loss of MeCP2 affects the expression of brain-derived neurotrophic factor (BDNF), an important neurotrophin which signals through its high affinity receptor tropomyosin receptor kinase B (TrkB, encoded by *Ntrk2* gene) [9–11]. *Mecp2-null* mice showed a significant downregulation of BDNF in CNS which was associated with developmental regression [11]. In addition, *Mecp2-null* mice and mice with loss of optimal BDNF – TrkB signaling in the CNS caused by a conditional loss of BDNF from post-mitotic neurons exhibited similar neuropathological phenotype [11]. These data implicate MeCP2-driven loss of BDNF – TrkB signaling as the mechanism driving cognitive and motor dysfunction associated with RTT. Here, we hypothesize that similar to its effect in CNS, MeCP2 loss alters BDNF expression in the ENS to cause GI dysmotility in the *Mecp2-null* mouse model of RTT.

BDNF – TrkB signaling in the ENS is significant for optimal neuronal activity [12]. Mice with reduced BDNF levels exhibit delayed GI transit, decreased stool frequency, and slow fecal pellet propulsion, and BDNF loss and mutations are associated with constipation and decreased stool frequency in human patients [13, 14]. We have recently identified the nature of *Bdnf* isoforms that are expressed in the post-natal ENS and found that *Bdnf isoforms IV and VI* (transcribed from exons 4 and 6, respectively) dominate the ENS [15]. Here, we use the *Mecp2-null* transgenic mice to assess whether GI motility regresses with age in these animals, and whether dysmotility is associated with specific changes to enteric *Bdnf* expression. Further, we assess whether *Mecp2-null* mice show significant changes to the overall ENS structure or loss of specific neuronal populations to provide a better understanding of the mechanisms driving GI dysmotility in RTT patients.

## Materials and Method

### Animals

All procedures were approved by the Institutional Animal Care and Use Committees (IACUC) at Beth Israel Deaconess Medical Center and Boston Children’s Hospital in accordance with the guidelines provided by the National Institutes of Health. Animals were maintained on a 12-hour light/dark cycle with ad libitum access to standard rodent chow (LabDiet, 5058) and water. *Mecp2*-deficient mouse line (B6.129P2(C)-*MeCP2*tm1.1Bird/J, JAX# mice:003890) heterozygous females were crossed with wild-type (WT) male C57BL/6 animals. Given the X-linked nature of the *Mecp2* gene, male mice carrying the *Mecp2*-mutant allele were effectively *Mecp2-null* genotype. All experiments were performed using male WT and *Mecp2-null* mice between 35 and 55 days of age. All control animals were male wild-type (WT) age-matched littermates of the mutant mice. Furthermore, we used the *Wnt1*-cre mice (C57BL/6 background) that were bred with the floxed TrkB.FL mice (STOCK Ntrk2tm1Ddg/J, Jax mice: 022363) to generate *Wnt1*-cre:*TrkB^fl/fl^*and *Wnt1*-cre:*TrkB^WT/WT^* mice that we have previously published on [15]. Specifically, 2-month-old male mice from both genotypes were used for this study.

### Whole gut transit time assay (WGTT) in mice

Whole gut transit time assay was performed as previously described [15]. Assay was performed only on those *Mecp2-null* mice that did not exhibit any visible signs of distress and discomfort. Briefly, Mice were individually caged in clean plastic cages and were left undisturbed for 30 minutes. These experiments were initiated between 8 and 9 AM. Water was provided to the mice during the experiment *ad libitum*. All mice received oral gavage of 0.3ml of 6% (w/v) carmine-red (Sigma C1022) in 0.5% (w/v) methylcellulose (Sigma M0512) in sterile saline. Seventy minutes after mice oral gavage, their individual cages were monitored every 15 minutes for the presence of red colored fecal pellets, and the time difference between the gavage and red fecal pellet production was measured as the whole gut transit time for that mouse. The experiment was terminated at 250 minutes post-gavage and the WGTT of any mice that did not expel the red dye at the termination was marked at the value of 265 (i.e. 250 + 15) min. A statistical analysis was performed on the mean difference in WGTT (in minutes) between groups.

### Mouse small intestine longitudinal muscle containing myenteric plexus (LM-MP) tissue isolation

LM-MP tissues contain the myenteric neurons and glial cells. This was performed only on those wildtype and *Mecp2-null* mice that did not exhibit any visible signs of distress and discomfort. To remove this tissue for analyses, mice were anesthetized with isoflurane and euthanized by cervical dislocation. Mice were placed in dorsal recumbency on the surgical surface and the abdominal skin was disinfected with 70% EtOH before performing a laproscopy to remove the small intestinal (SI) tissue in a petri dish containing sterile 1X PBS. Small intestine was flushed with the sterile 1X PBS solution using a 20ml syringe to clear luminal contents. Small intestine was cut into 2 cm segments, then kept in a sterile petri dish containing sterile Opti-MEM medium. Each SI segment was placed over a sterile 1 ml pipette and LM-MP tissue was peeled off from the underlying tissue using a wet sterile cotton swab. Tissues isolated for quantitative real time PCR (qRT-PCR)-based assessments were snap frozen immediately after isolation. Tissues isolated for immunostaining experiments were flattened, fixed in freshly prepared ice-cold 4% paraformaldehyde (PFA) solution for 5 minutes based on our established protocol [16] and then stored in 1X sterile PBS for subsequent immunostaining procedures.

### Cell culture experiments

Neurospheres were generated from cells isolated from the small intestine of wild-type (WT) and/or *Mecp2* knockout (KO) male mice at postnatal day 35 (P35), as previously described [17]. Briefly, the longitudinal muscle–myenteric plexus (LM-MP) layer was dissected from the small intestine and subjected to enzymatic dissociation in M199 medium supplemented with 0.1% BSA, Trehalose, 20 mM HEPES, 50 U/mL DNAse I, and 1.1 mg/mL collagenase (Worthington). Tissue digestion was performed at 37°C for 40 min. The resulting cell suspension was centrifuged at 210g for 7min, washed once with PBS, and resuspended in stem cell medium (SCM). The suspension was passed through a 100-µm cell strainer to obtain a single cell suspension. Cells were seeded in T25 flasks (corning) and maintained in SCM consisting of Neurobasal medium supplemented with B27 supplement, 0.7% BSA, 2 mM L-glutamine, 100 U/ml antibiotic–antimycotic solution, and 20 ng/ml each of fibroblast growth factor (FGF), epidermal growth factor (EGF), and glial cell line-derived neurotrophic factor (GDNF). Cultures were maintained under standard cell culture conditions. Half of the culture medium was replaced every 48h with fresh SCM supplemented with growth factors. After 1 week of culture, primary neurospheres were dissociated using Accutase and replated onto fibronectin-coated (100ug/ml) 12-well culture plates for differentiation and pharmacological treatment.

## Experiments on differentiated enteric neurons

Three different combinations of experiments were performed on differentiating cells derived from neurospheres. In the first experiment *E1*, differentiating cells from WT mice (n = 4/group, males) were differentiated in SCM with Vehicle (Control) or with TrkB antagonist ANA-12 (10 nM, Tocris Biosci) for 14 days. In the second experiment *E2*, differentiating cells from wildtype (WT) C57BL/6 male mice and *Mecp2* KO mice (n = 3/group, males) were differentiated in SCM without any growth factors for 14 days. In the third experiment *E3*, differentiating cells from *Mecp2* KO mice (n = 3/treatment; males) were cultured in SCM with Vehicle (Control) or with TrkB agonist HIOC (0.5 µM, Tocris Biosci) for 7 days. Media in the cultures was changed every alternate day. At the end of the culture period, cells were lysed and total RNA was isolated using Direct-zol RNA miniprep kit (Zymo research), and gene expression was quantified by quantitative real time PCR (qRT-PCR) using gene specific primers.

### Quantitative real-time Polymerase Chain Reaction (qRT-PCR)

Total RNA was extracted from cells and from frozen LM-MP tissues using Direct-zol RNA miniprep kit (Zymo research) and 1 μg RNA was reverse transcribed to cDNA using iScript cDNA synthesis kit (Biorad, Cat.No.1708891) in a 20μl reaction volume. The qRT-PCR assay was performed with TaqMan Fast advanced or powerup SYBR qPCR Master Mix (Applied biosystems) on a QuantStudio3 real time PCR system using gene specific primers listed in **Table 1**. A schematic of TrkB.FL and TrkB.T1 isoform-specific primers is provided in **Supplementary Fig 1**. Mouse *Hprt* served as a housekeeping gene for normalization. 20μl qRT-PCR reactions were incubated at 95℃ for 10 min, followed by 40 cycles at 95℃ for 10 s, 60℃ for 10 s, and 72℃ for 40 s. Relative fold changes of genes between groups were calculated using the 2^−ΔΔCt^ method.

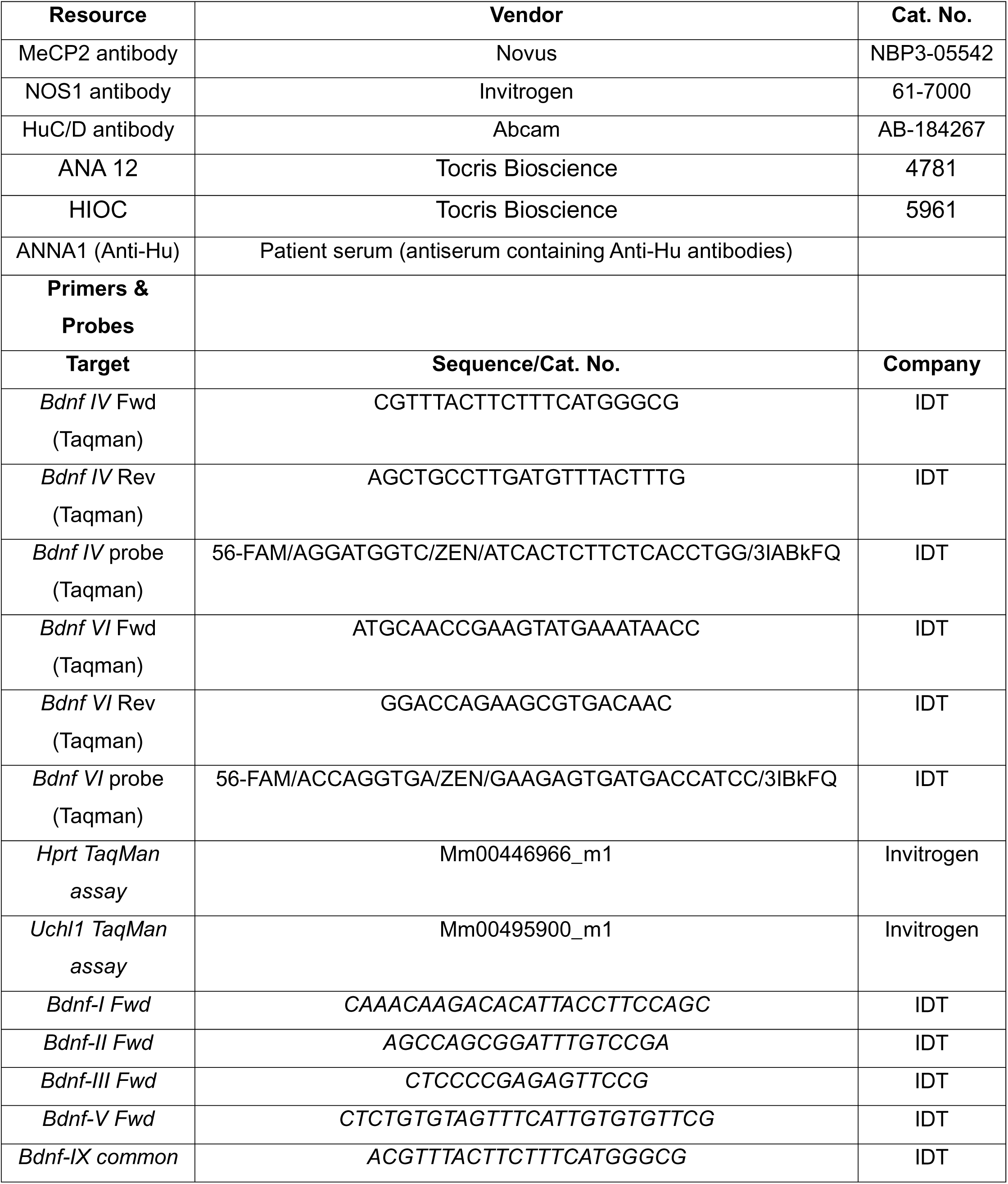
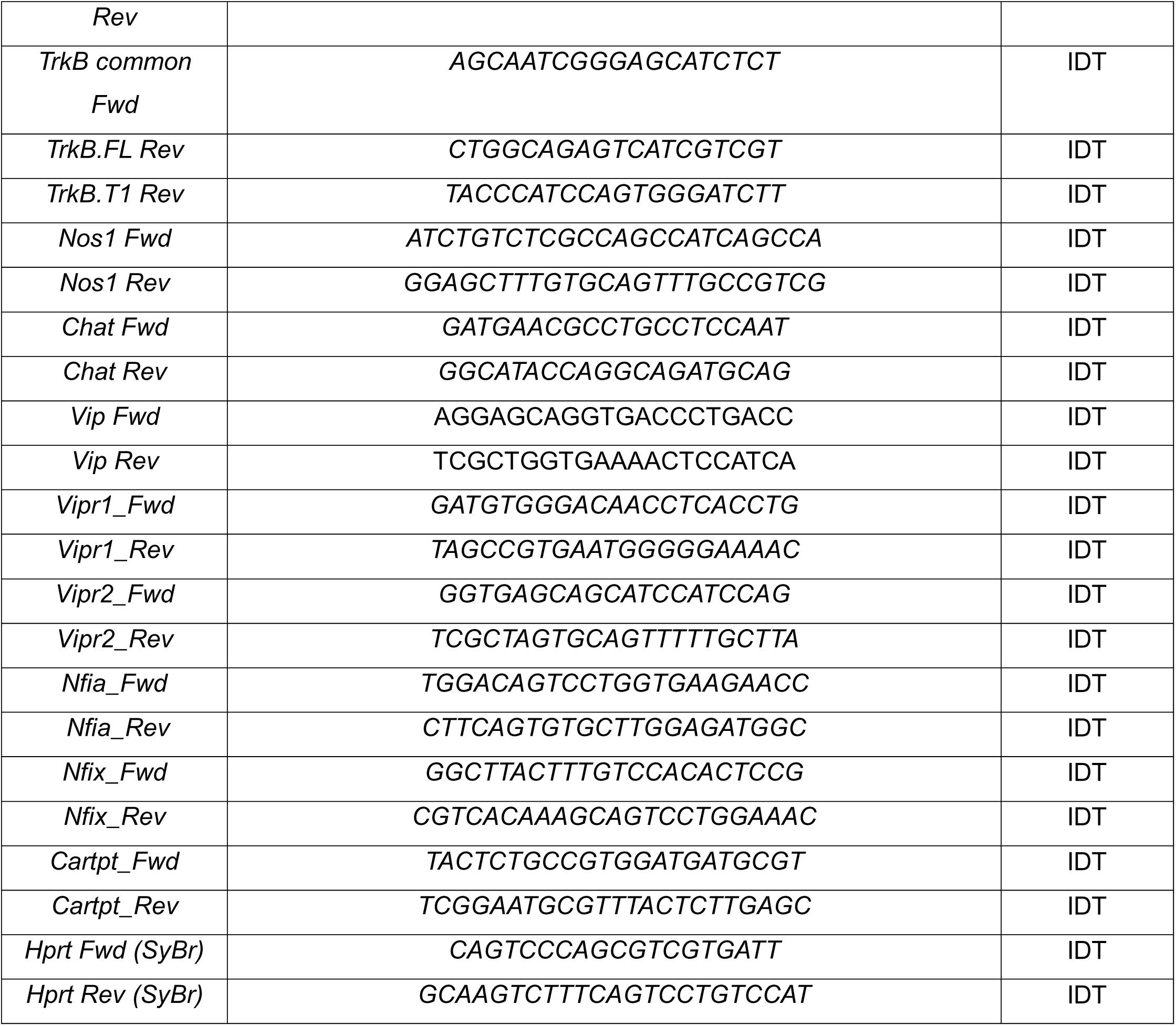
Table. 1.

### Immunostaining

Fixed longitudinal muscle containing myenteric plexus (LM-MP) tissues were blocked and permeabilized with blocking buffer contain 5% normal goat serum and 0.5% Triton X-100 in 1X sterile PBS for 60 minutes at room temperature. For assessing the expression of MeCP2 in myenteric neurons, we performed immunostaining of adult C57BL/6 and *Mecp2-null* mice with anti-MeCP2 antibody (1:500) and ANNA-1 patient serum containing anti-Hu antibodies (1:1000). After counterstaining with anti-Rabbit conjugated with Alexa fluor-647 and anti-human conjugated with Alexa fluor-488 antibodies and nuclear dye DAPI, the tissues were mounted with Prolong Gold anti-fade mountant (Invitrogen) and imaged under Leica Stellaris Confocal microscope using 63X PL APO oil immersion objective (NA: 1.40) with z-step set at 0.85 μM.

To evaluate neuronal structure and nNOS-positive neurons, tissues were incubated either with rabbit anti-Hu (1:500 in 0.1% Triton X-100) or with rabbit anti-nNOS (1:250 in 0.1% Triton X-100) primary antibodies. For secondary staining, Alexa Fluor 488–conjugated anti-rabbit antibodies were used, along with the nuclear dye DAPI imaged with a fluorescence microscope (EVOS). For each tissue, at least 10 random fields were acquired using a 40X objective.

### Single nucleus/cell RNA sequencing data analyses

We data-mined publicly available single cell/nucleus RNA sequencing data compilation, generated by combining data from juvenile and adult ENS neurons by the Southard-Smith lab for this study [18]. The data compilation used has been made publicly by the Southard-Smith lab at https://zenodo.org/records/17420912. Graphs were made using UMAP representation of the data using Cellxgene.

### Statistical analysis

GraphPad Prism version 10.0 (GraphPad Software, San Diego, CA, USA) was used for graph generation, and statistical evaluation. Data, expressed as the mean ± standard error of the mean (SEM), were analyzed using t-test. differences were considered statistically significant at p < 0.05.

## Results

### MeCP2 is expressed by murine ENS cells including myenteric neurons of the adult small intestinal tissue

Confocal microscopy of small intestinal LM-MP from adult wildtype mice immunostained with antibodies against MeCP2 and pan-neuronal marker Hu shows that MeCP2 is present as punctate bodies in the nucleus in enteric neurons as well as in other cells of the myenteric ganglia. We confirmed the specificity of this antibody by observing that similar immunohistochemistry using the small intestinal LM-MP tissue from P55 *Mecp2-null* male mouse showed a lack of MeCP2 immunostaining in all cells (**Fig 1**)

**Figure 1:**
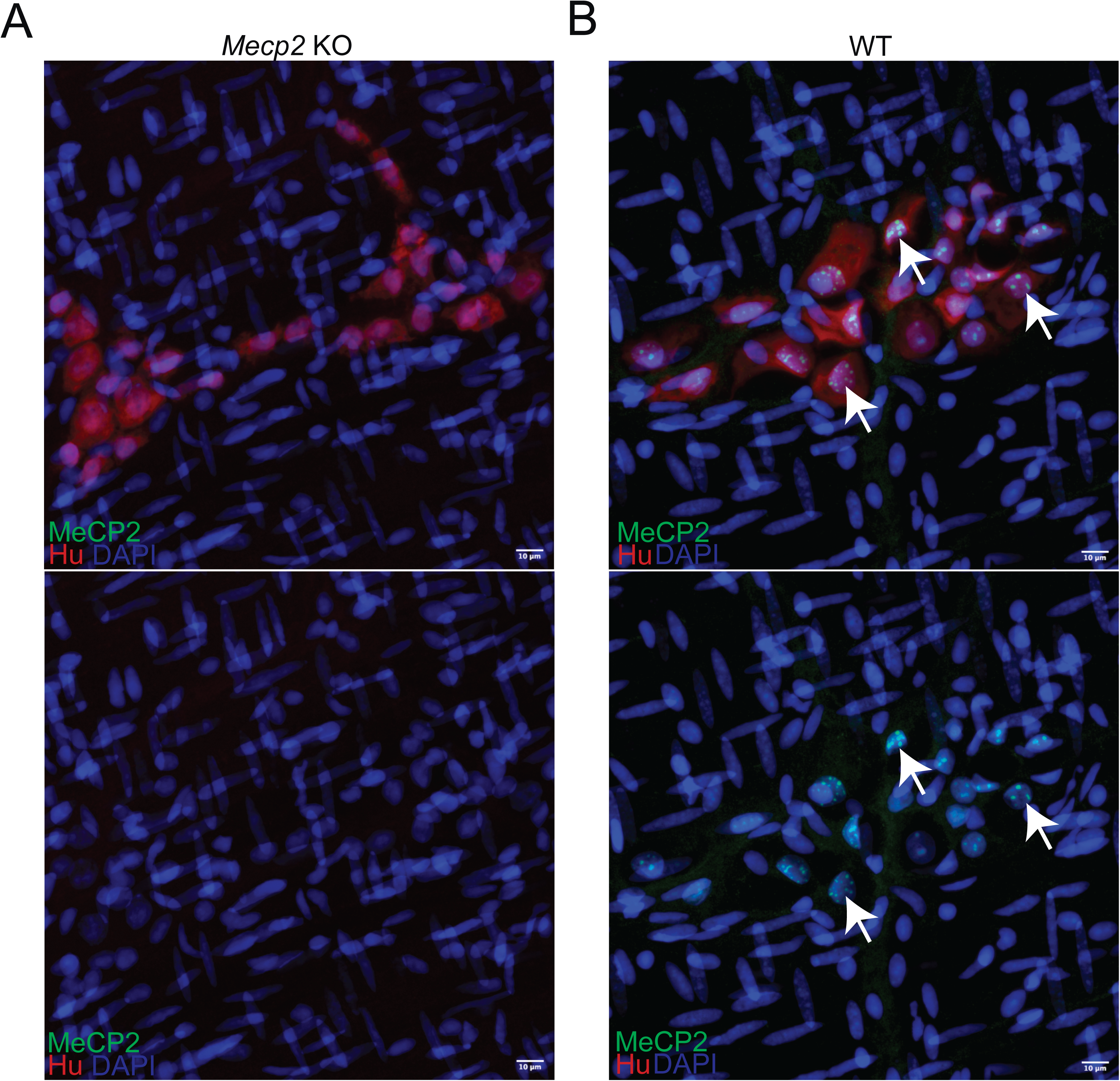
Nuclear MeCP2 is detected in enteric neurons and surrounding cells of the myenteric ganglia in wildtype but not in *Mecp2-null* mice. Representative images of the adult small intestinal longitudinal muscle – myenteric plexus (LM-MP) preparations from (A) P55-aged *Mecp2-null*; and (B) wildtype (WT) mice, when immunostained with antibody against MeCP2 (green) and ANNA1 patient-derived serum containing anti-Hu antibodies (red) and counterstained with nuclear dye DAPI (blue) shows that MeCP2 protein is detected within the nucleus of Hu-immunolabeled cells of the myenteric ganglia of WT mice (white arrows), but not of the *Mecp2-null* mice. Scale bar = 10 µm.

### Loss of MeCP2 protein impairs whole GI motility in *Mecp2-null* male mice as they near adulthood

We next investigated whether the loss of MeCP2 causes GI dysmotility before or after the onset of developmental regression. For this, we utilized male mice of two ages: post-natal day 35 (Juvenile, P35) and post-natal day 55 (near adulthood, P55), and experiments were performed between the *Mecp2-null* mice and littermate age-matched sex-matched wildtype (WT) control mice at the two aforementioned ages. By performing carmine red dye-based whole gut transit time (WGTT) experiments, we found that *Mecp2-null* mice at P35 age (juvenile stage) did not show any significant loss of optimal GI motility when compared to WT mice (n = 5/group; mean ± S.E. of WGTT (in min): WT: 118.0 ± 18.61; *Mecp2-null*: 95.00 ± 8.36, p = 0.29, Students’ t-test), whereas at P55 age (near adulthood) *Mecp2-null* mice exhibited significantly reduced GI motility when compared to age-matched WT male mice (n = 6/group; mean ± S.E. of WGTT (in min): WT: 125.0 ± 18.44; *Mecp2-null*: 217.50 ± 24.92, p = 0.013, Students’ t-test; **Fig 2**). This shows that GI motility of *Mecp2-null* mice undergoes functional regression as they chronologically age from adolescence to near-adulthood.

**Figure 2:**
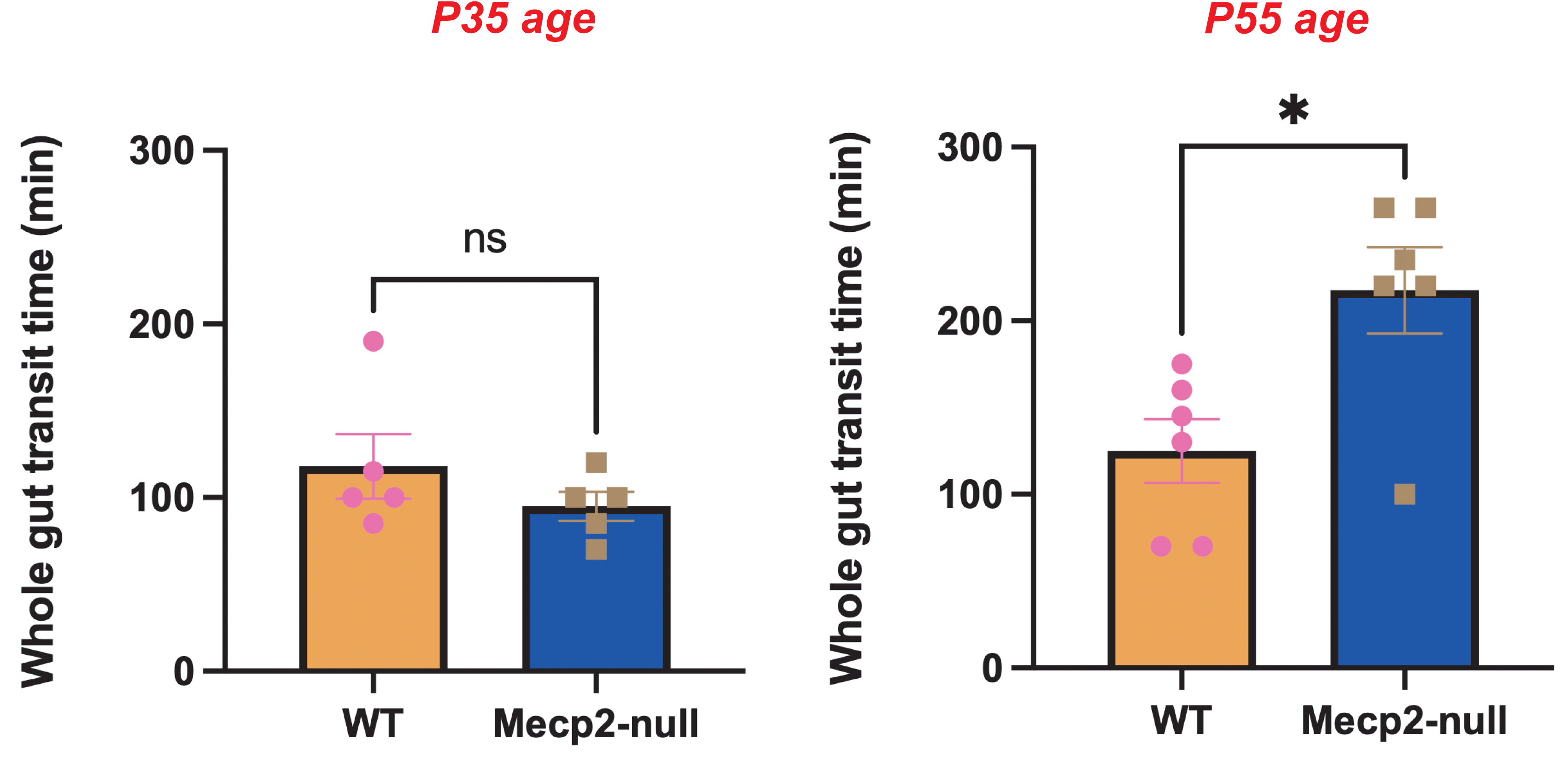
MeCP2 loss drives maturation-associated loss of optimal whole GI motility. Whole GI transit time (WGTT) analyses to assess the time taken for orally gavaged carmine red dye to traverse the entire gastrointestinal tract and show in feces (in minutes) was performed in *Mecp2-null* male mice and age-and sex-matched wildtype (WT) littermate mice at two time points of post-natal day 35 (P35) and post-natal day 55 (P55). The two genotypes did not show any significant difference in their WGTT at the P35 age, but the *Mecp2-null* mice showed significantly increased WGTT at the P55 age, when compared to WT control mice (* p < 0.05; Students’ t-test).

### Specific *Bdnf* transcripts are reduced in the small intestinal LM-MP of *Mecp2-null* mice, without significant differences in TrkB isoform transcript expression

We investigated whether specific *Bdnf* transcripts were differentially expressed in the small intestinal LM-MP of *Mecp2-null* and WT mice. Given that we have found that *Bdnf IV* (transcription initiation from exon 4) and *Bdnf VI* (transcription initiation from exon 6) together account for more than 90% of *Bdnf* isoforms in the post-natal murine small intestinal ENS [15], we tested how the expression of these isoforms differs between P55 WT and *Mecp2-null* male mice, which we observed to have significant GI dysmotility (**Fig 3**). Using qRT-PCR with primers specific to *Bdnf IV* and *Bdnf VI* isoforms, we found significantly lower expression of both *Bdnf IV* and *Bdnf VI* transcripts in small intestinal LM-MP from P55 *Mecp2-null* males compared with WT controls (n = 6/WT and 4/*Mecp2-null*; mean ± S.E. fold change of *Bdnf IV*: WT: 1.015 ± 0.076; *Mecp2-null*: 0.170 ± 0.043, p < 0.0001; mean ± S.E. fold change of *Bdnf VI*: WT: 1.003 ± 0.034; *Mecp2-null*: 0.654 ± 0.026, p < 0.0001; Students’ t-test; **Fig 3A, B**).

**Figure 3:**
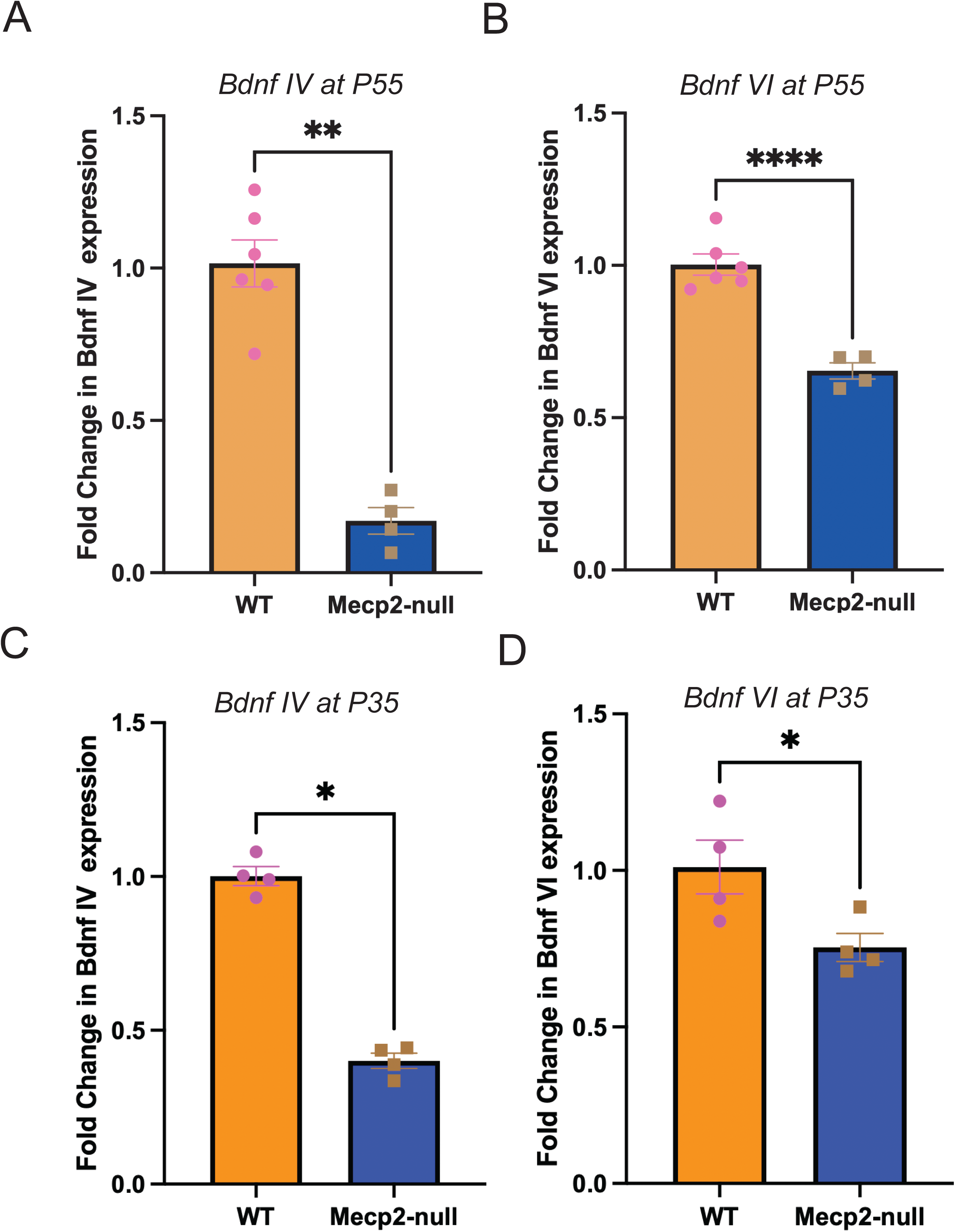
*Bdnf IV* and *VI* transcript expression is reduced in small intestinal LM-MP from *Mecp2-null* males. Quantitative real time PCR (qRT-PCR) analyses of *Bdnf IV* and *Bdnf VI* isoforms in the small intestinal longitudinal muscle – myenteric plexus (LM-MP) tissues of (A, B) P55 mice and in (C, D) P35-aged male *Mecp2-null* and age-and sex-matched wildtype (WT) littermate mice show that the relative abundance (fold change) of these two isoforms is significantly reduced in *Mecp2-null* mice compared to WT controls (*p < 0.05, ** p < 0.01, **** p < 0.001; Students’ t-test).

We next compared the expression of less abundant *Bdnf* transcripts between genotypes. Expression of *Bdnf II* was significantly lower in *Mecp2-null* mice (n = 4/group mean ± S.E. fold change of *Bdnf II*: WT: 1.061 ± 0.180; *Mecp2-null*: 0.092 ± 0.047, p = 0.002; Students’ t-test), whereas no statistically significant differences were detected for *Bdnf I, III*, or *V* isoforms (n = 4/group, mean ± S.E. fold change of *Bdnf I*: WT: 1.183 ± 0.383; *Mecp2-null*: 0.671 ± 0.216, p = 0.289; mean ± S.E. fold change of *Bdnf III*: WT: 2.132 ± 0.892; *Mecp2-null*: 0.717 ± 0.430, p = 0.203; mean ± S.E. fold change of *Bdnf V*: WT: 1.748 ± 0.819; *Mecp2-null*: 1.794 ± 0.625, p = 0.965; Students’ t-test).

BDNF mediates its downstream effects by binding to high-affinity TrkB receptor, encoded by the *Ntrk2* gene. Alternative splicing of *Ntrk2* generates two primary isoforms: the full-length receptor (TrkB.FL), which initiates intracellular signaling cascades, and a truncated receptor (TrkB.T1), which lacks a kinase domain and functions as a competitive “sink” for BDNF [19]. Since we have observed that BDNF and TrkB are expressed by distinct neuronal populations in the ENS [15], we next compared expression of the two TrkB variants between P55 *Mecp2-null* and WT males. qRT-PCR analyses using small intestinal LM-MP tissues from P55-aged male *Mecp2-null* and age-and sex-matched littermate wildtype (WT) mice revealed that expression of neither TrkB.FL nor TrkB.T1 showed statistically significant differences between the two genotypes (n = 4/group, mean ± S.E. fold change of TrkB.FL: WT: 1.020 ± 0.112; *Mecp2-null*: 0.986 ± 0.199, p = 0.887; mean ± S.E. fold change of TrkB.T1: WT: 1.040 ± 0.181; *Mecp2-null*: 3.919 ± 1.535, p = 0.112; Students’ t-test).

We next examined whether reduced expression of the dominant enteric *Bdnf IV* and *VI* transcripts was also detectable at P35, when WGTT did not differ significantly between genotypes. We found that these two *Bdnf* isoforms were also significantly reduced at the P35 age (n = 4/group, mean ± S.E. fold change of *Bdnf IV*: WT: 1.001 ± 0.030; *Mecp2-null*: 0.400 ± 0.024, p < 0.0001; mean ± S.E. fold change of *Bdnf VI*: WT: 1.011 ± 0.085; *Mecp2-null*: 0.754 ± 0.044, p = 0.03; Students’ t-test; **Fig 3C, D**). Neither TrkB.FL nor TrkB.T1 showed significant change in gene expression at this age (n = 4/group, mean ± S.E. fold change of TrkB.FL: WT: 1.029 ± 0.136; *Mecp2-null*: 0.975 ± 0.093, p = 0.759; mean ± S.E. fold change of TrkB.T1: WT: 1.113 ± 0.289; *Mecp2-null*: 1.546 ± 0.390, p = 0.407; Students’ t-test). Reduced *Bdnf IV* and *VI* transcript expression at both ages, without significant differences in TrkB isoform transcripts, is consistent with potentially reduced BDNF availability.

### Loss of MeCP2 does not alter overall neuronal numbers or abundance of nitrergic neurons in the small intestinal myenteric plexus

GI dysmotility has often been linked to alterations in the total numbers of neurons, where both a reduction and increase in total neuronal numbers is known to drive GI dysmotility [17, 20]. Thus, it is important to assess whether loss of MeCP2 led to a significant change in the overall neuronal numbers in the ENS. We performed immunohistochemical analysis of small intestinal LM-MP tissues from P55-aged male *Mecp2-null* and age-and sex-matched littermate wildtype (control) mice with antibodies against pan-neuronal marker Hu. Quantification of Hu-immunolabeled cells per 40X field of view showed comparable neuronal densities in WT and *Mecp2-null* mice (n = 3/group: mean ± S.E. of Hu-immunolabeled cells per 40X field: WT: 34.80 ± 1.59; *Mecp2-null*: 35.32 ± 3.14; p = 0.89, Students’ t-test; **Fig 4A**).

**Figure 4:**
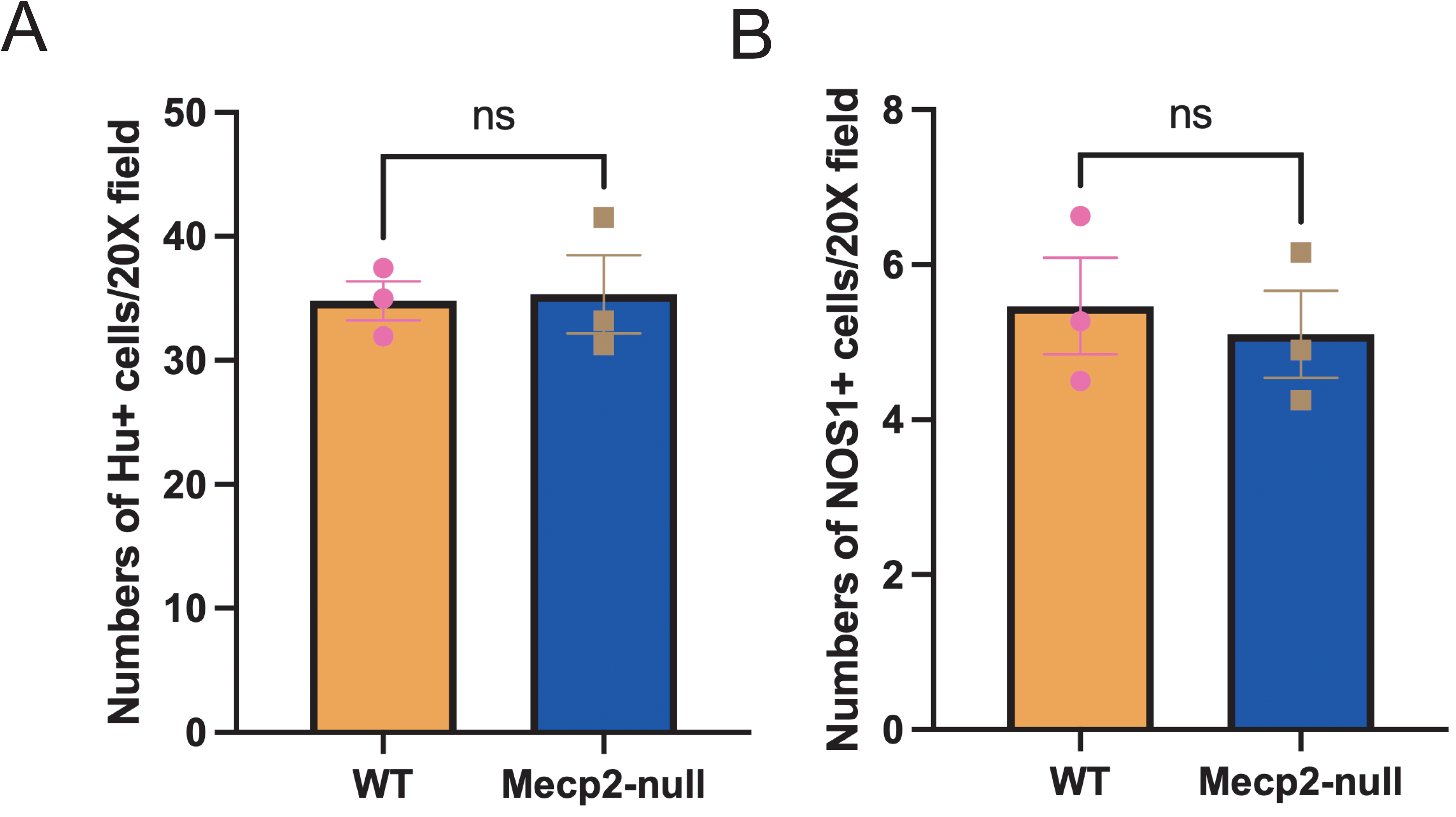
Loss of MeCP2 does not alter abundance of mature neurons or that of nitrergic neurons in the small intestinal myenteric plexus of dysmotile male mice. P55-aged male *Mecp2-null* and age-and sex-matched wildtype (WT) littermate mice show no significant differences in the numbers of (A) Hu-immunolabeled neurons enumerated in 40X field or in the (B) numbers of NOS1-immunolabeled nitrergic neurons enumerated in 40X field (Students’ t-test).

Since a prior report suggested alterations in the expression of NOS1 protein in the myenteric plexus of *Mecp2-null* mice, we next tested whether these mice show changes in the abundance of NOS1-immunolabeled nitrergic neurons, which function as inhibitory neurons in the small intestinal myenteric plexus. For this, we performed immunohistochemical analysis of small intestinal LM-MP tissues from P55-aged male *Mecp2-null* and age– and sex-matched littermate wildtype (control) mice with antibodies against NOS1. Quantification of NOS1-immunolabeled cells per 40X field of view showed no statistically significant differences in the abundance of nitrergic neurons between WT and *Mecp2-null* mice (n = 3/group: mean ± S.E. of Hu-immunolabeled cells per 40X field: WT: 5.46 ± 0.62; *Mecp2-null*: 5.10 ± 0.56; p = 0.68, Students’ t-test; **Fig 4B**).

### Loss of MeCP2 does not change expression of *Nos1* and *Chat* genes but does significantly alter expression of *Vip* in the small intestinal myenteric plexus

It is plausible that while MeCP2 loss does not change abundance of nitrergic neurons, it may alter the expression of *Nos1* or the expression of gene encoding Choline acetyltransferase (*Chat*) which is expressed by acetylcholine releasing excitatory neurons in the myenteric plexus. qRT-PCR analyses using primers specific to *Nos1* or *Chat* showed no significant differences in the expression of these genes between P55-aged *Mecp2-null* male mice and age– and sex-matched wildtype (WT) littermate control mice (n = 4/group, mean ± S.E. fold change of *Nos1*: WT: 1.133 ± 0.367; *Mecp2-null*: 1.486 ± 0.162, p = 0.41; mean ± S.E. fold change of *Chat*: WT: 1.079 ± 0.232; *Mecp2-null*: 1.780 ± 0.331, p = 0.13; Student’s t-test; **Fig 5A, B**). Consistent with the lack of change in overall neuronal numbers, we tested and found that the expression of the pan-neuronal gene Uchl1 (encoding PGP9.5 protein expressed by all enteric neurons) was not significantly altered between the two genotypes (n = 4/group, mean ± S.E. fold change of *Uchl1*: WT: 1.027 ± 0.128; *Mecp2-null*: 1.153 ± 0.239, p = 0.65; Student’s t-test; **Fig 5C**).

**Figure 5:**
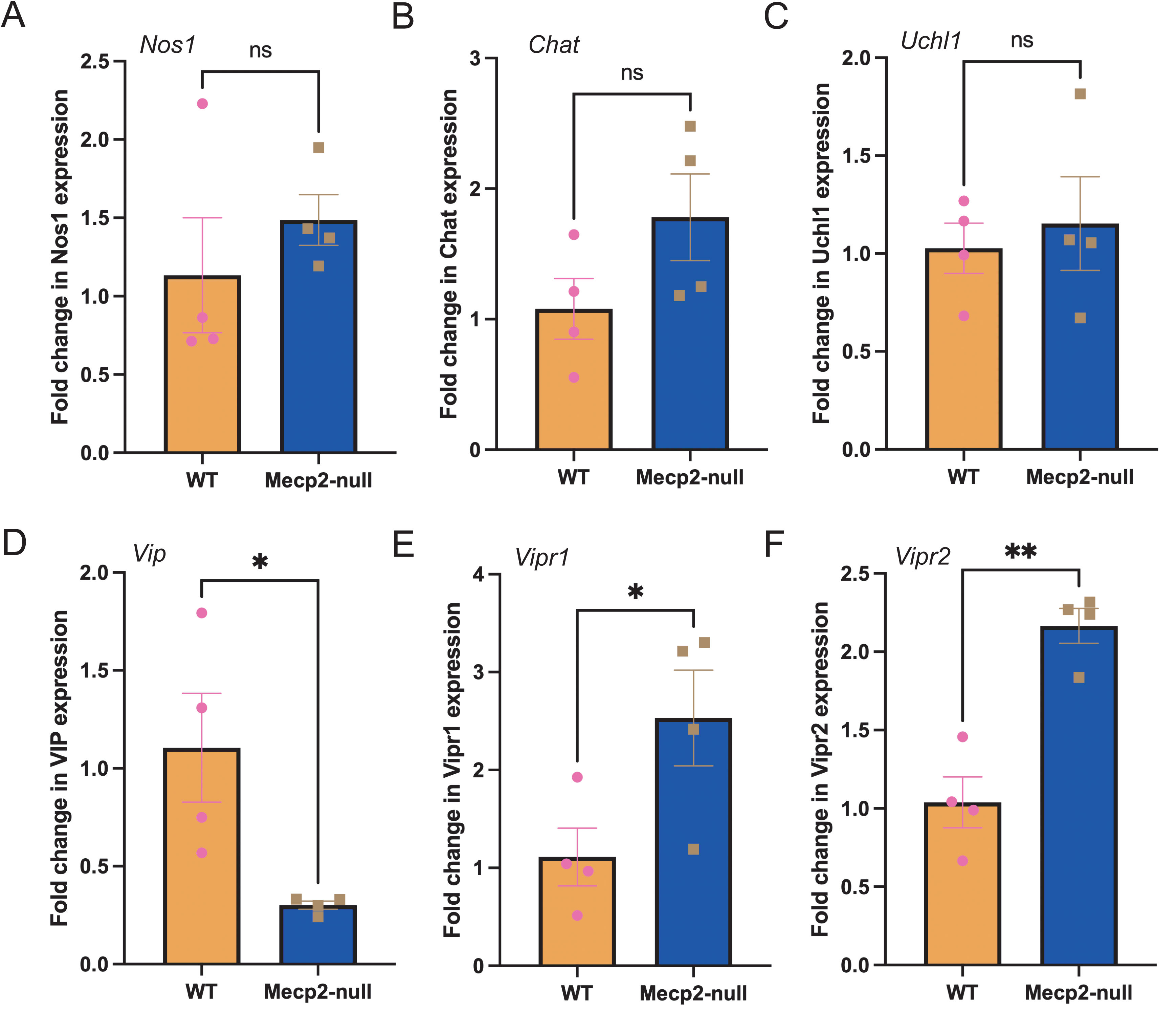
Loss of MeCP2 alters VIPergic signaling pathway genes in the small intestinal myenteric plexus of P55 dysmotile male mice. P55-aged male *Mecp2-null* and age-and sex-matched wildtype (WT) littermate mice show no significant differences in the relative abundance of (A) nitrergic neuronal marker gene Nos1 (B) cholinergic marker gene *Chat*, (C) or the pan-neuronal marker gene *Uchl1* coding the protein PGP9.5 in the small intestinal myenteric plexus tissue (Students’ t-test). However, P55-aged male *Mecp2-null* mice show (D) significant reduction in the expression of inhibitory neuropeptide gene *Vip*, and a significant increase in the expression of VIP receptor genes (E) *Vipr1* encoding VPAC1R and (F) *Vipr2* encoding VPAC2R in the small intestinal LM-MP tissues (* p < 0.05, ** p < 0.01; Students’ t-test).

The central circadian clock (suprachiasmatic nucleus) in *Mecp2-null* mice exhibited significant reduction in the numbers of VIP-expressing neurons and an associated alteration in circadian rhythm [21]. Approximately 25% of all myenteric neurons in the small intestine express the inhibitory neuropeptide VIP (*Vip*), and VIPergic signaling in the gut has been shown to be a significant driver of GI motility [22–26]. The small intestinal myenteric neurons are known to express receptors for VIP (*Vipr1* and *Vipr2*) [27, 28]. Hence, we decided to test whether loss of MeCP2 alters the expression of *Vip, Vipr1*, and *Vipr2* in the small intestinal myenteric plexus. We found that P55-aged *Mecp2-null* mice indeed showed a significant reduction in the expression of *Vip* (n = 4/group, mean ± S.E. fold change of *Vip*: WT: 1.105 ± 0.278; *Mecp2-null*: 0.301 ± 0.021, p = 0.028; Student’s t-test; **Fig 5D**), but a significant increase in the expression of the VIP receptor genes (n = 4/group, mean ± S.E. fold change of *Vipr1*: WT: 1.11 ± 0.29; *Mecp2-null*: 2.53 ± 0.49, p = 0.047; mean ± S.E. fold change of *Vipr2*: WT: 1.03 ± 0.16; *Mecp2-null*: 2.16 ± 0.11, p = 0.001; Student’s t-test; **Fig 5 E,F**). These data show that loss of MeCP2 affects VIPergic signaling in the small intestinal LM-MP tissue in dysmotile mice at the P55 age.

Given that significant reduction in *Bdnf IV* and *VI* levels was observed at both P35 and P55 ages, while delayed WGTT was observed at P55 age, we tested whether the expression of *Vip, Vipr1* and *Vipr2* was significantly altered only at the P55 dysmotile age, or at both ages. qRT-PCR analyses showed that P35 mice showing no significant alteration in WGTT do now show any significant difference in the expression of *Vip,* or *Vipr1*, but show a strong tendency of increase in expression of *Vipr2* gene (n = 4/group, mean ± S.E. fold change of *Vip*: WT: 1.179 ± 0.303; *Mecp2*-null: 0.897 ± 0.057, p = 0.397; mean ± S.E. fold change of *Vipr1*: WT: 1.004 ± 0.050; *Mecp2*-null: 2.558 ± 0.856, p = 0.120; mean ± S.E. fold change of *Vipr2*: WT: 1.002 ± 0.033; *Mecp2*-null: 1.337 ± 0.137, p = 0.055; Student’s t-test; **Fig 6**). Across the two ages examined, reduced *Bdnf IV* and *VI* transcript expression was detectable at P35, whereas significant *Vip* downregulation and delayed transit were detected at P55, thus establishing an age-associated sequence.

**Figure 6:**
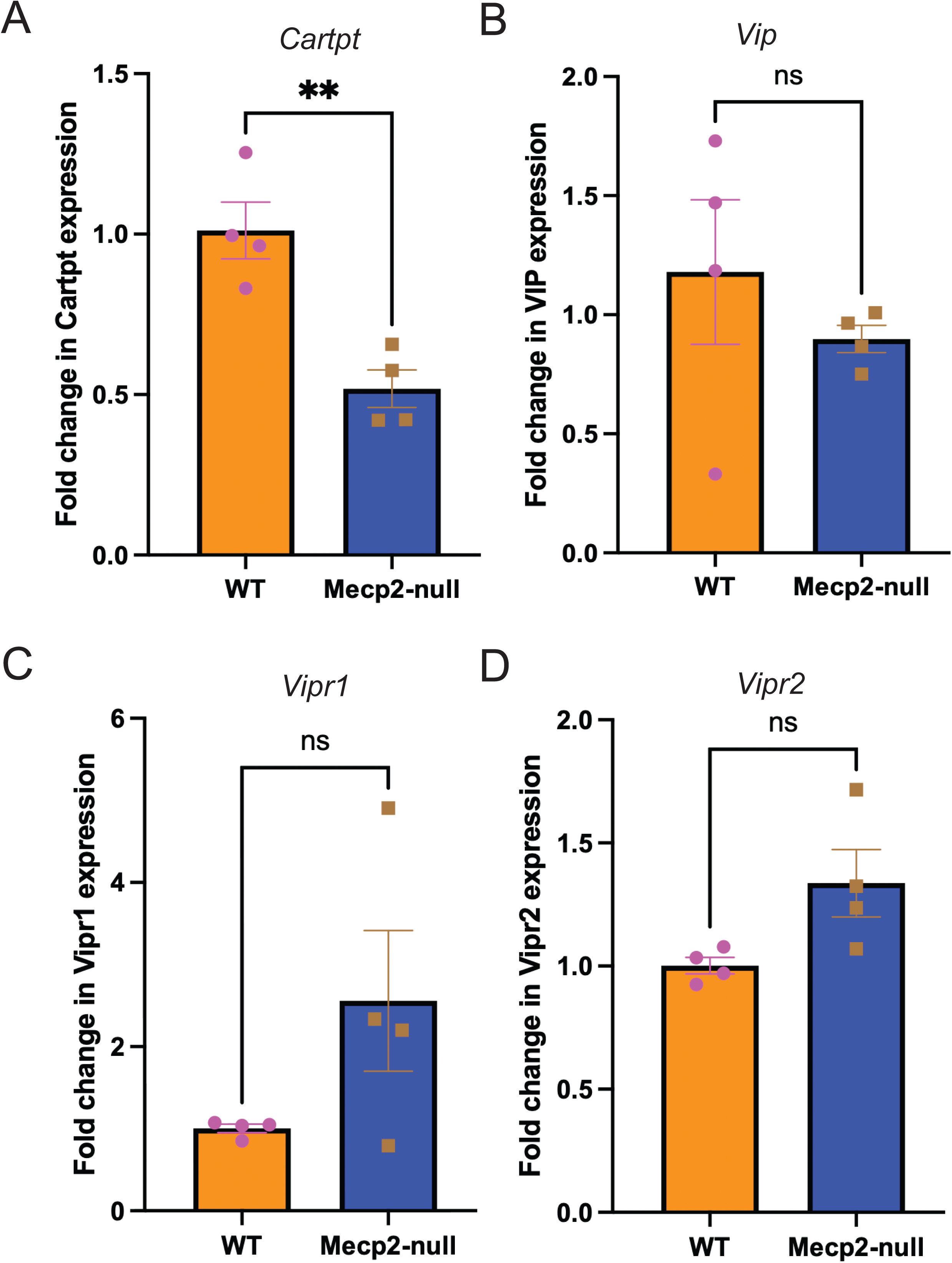
Expression of VIPergic signaling pathway genes is not significantly altered in the small intestinal myenteric plexus of P35 non-dysmotile male mice. P35-aged male *Mecp2-null* and age-and sex-matched wildtype (WT) littermate mice show (A) significant difference in the relative abundance of *Cartpt* transcripts; but not of (B) *Vip*, (C) *Vipr1*, and (D) *Vipr2* in the small intestinal LM-MP tissues (** p < 0.01; ns = p > 0.05; Students’ t-test).

*Vip*-expressing inhibitory neurons are a sub-population of inhibitory nitrergic neurons [29]. We mined publicly available single nucleus RNA seq data to observe that the cluster of inihibtory *Nos1*-expressing nitrergic neurons (**Fig 7A**) is subdivided in 3 sub-populations: Population A: comprising *Nos1^+^ Vip^low^* neurons, Population B comprising *Nos1^+^ Nfia^+^ Nfix^+^ Vip^-^*neurons, and finally Population C comprising *Nos1^+^ Nfia^-^Nfix^-^Vip^+^ Cartpt^+^* neurons (**Fig 7 B-F**). Significant reduction in *Vip* expression in *Mecp2-null* small intestinal myenteric plexus could suggest either reduced expression of *Vip*, or the loss of a specific subpopulation of *Vip^+^* neurons. We hypothesized that congenital loss of MeCP2 drives the loss of a specific VIPergic sub-population of neurons. We reasoned that loss of *Vip* and *Cartpt* expression together in *Mecp2-null* small intestinal ENS would suggest a loss of Population C, and the loss of transcription factors *Nfia* and *Nfix,* but not of *Cartpt,* would signify a loss of Population B nitrergic neurons. We performed qRT-PCR using primers specific to *Nfia, Nfix*, and *Cartpt* and found a significant reduction in the expression of *Cartpt* and a significant increase in the expression of *Nfia* (n = 4/group, mean ± S.E. fold change of *Cartpt*: WT: 1.00 ± 0.02; *Mecp2-null*: 0.37 ± 0.02, p < 0.0001; mean ± S.E. fold change of *Nfia*: WT: 1.02 ± 0.12; *Mecp2-null*: 1.63 ± 0.06, p = 0.005; Student’s t-test; **Fig 7 G,H**). Expression of *Nfix* did not show any statistically significant change compared to WT controls (n = 4/group, mean ± S.E. fold change: WT: 1.09 ± 0.21; *Mecp2-null*: 1.58 ± 0.12, p = 0.10; Student’s t-test; **Fig 7 I**). These data suggest that the relative abundance of *Vip^+^ Cartpt^+^ Nos1^+^* population C is significantly reduced, while that of *Nfia^+^ Nos1^+^* neurons (population B) is significantly increased in the P55-aged *Mecp2-null* male mice.

**Figure 7:**
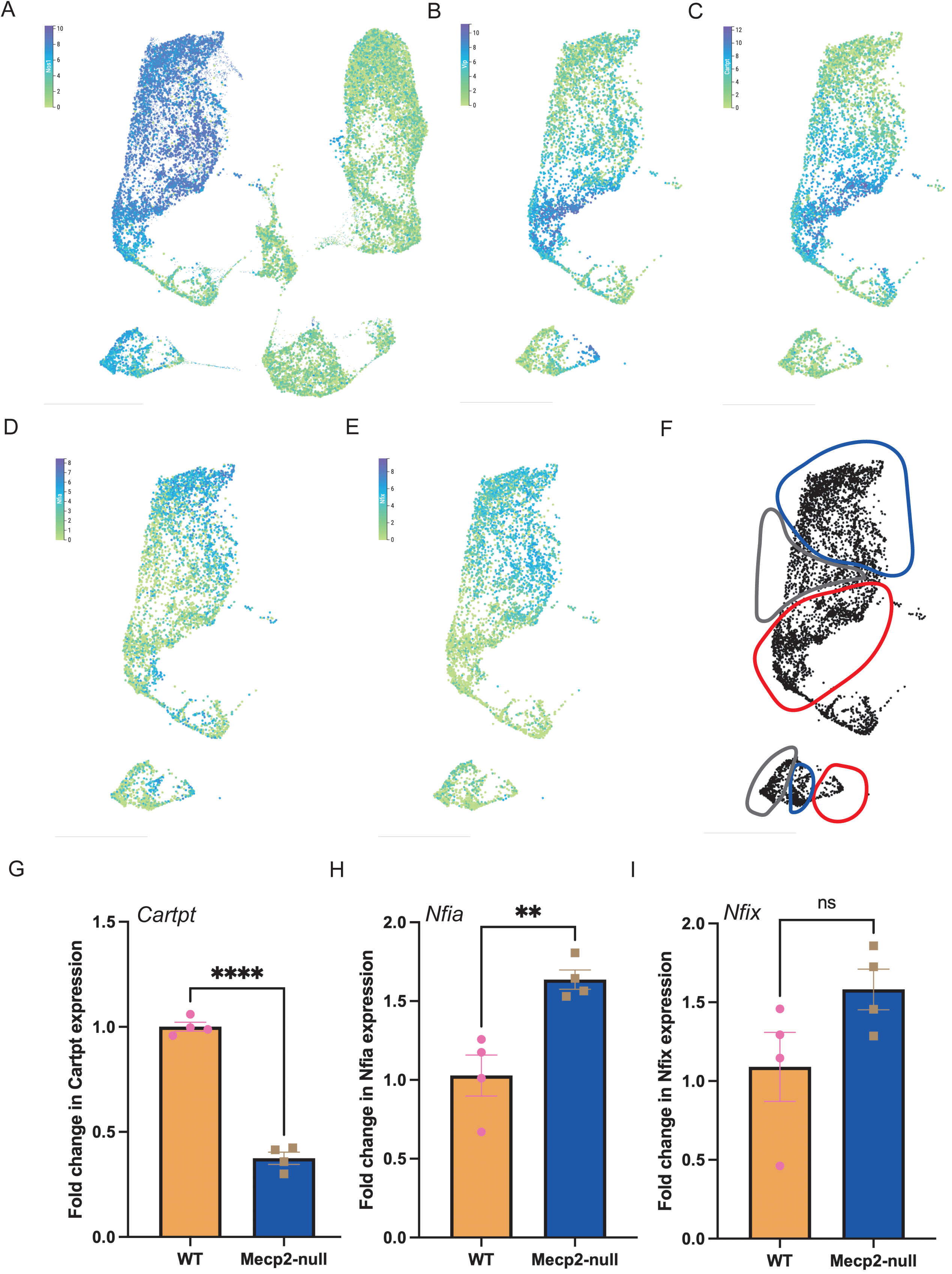
Loss of MeCP2 in small intestinal ENS drives putative changes in neuronal subpopulations of inhibitory myenteric neurons. Analyses of publicly available data shows (A) a UMAP representation of juvenile and adult enteric neuronal single cell/nucleus RNA sequencing data, where their expression of the inhibitory neuronal marker *Nos1* is queried to define the entire inhibitory neuronal population. The selected inhibitory neuronal population is then extracted and the expression of important sub-population level markers, namely (B) *Vip*, (C) *Cartpt,* (D) *Nfia,* and (E) *Nfix* is queried to show that *Vip* and *Cartpt* are expressed by the same inhibitory subpopulation, while *Nfia* and *Nfix* are expressed by a distinct inhibitory subpopulation of neurons. (F) Based on these marker expression profiles, we delineate three subpopulations of nitrergic neurons, namely Population A delineated in grey: comprising *Nos1^+^ Vip^low^* neurons, Population B delineated in Blue: comprising *Nos1^+^ Nfia^+^ Nfix^+^ Vip^-^*neurons, and finally Population C delineated in Red: comprising *Nos1^+^ Nfia^-^Nfix^-^Vip^+^ Cartpt^+^* neurons. qRT-PCR results show that the small intestinal LM-MP tissue of P55-aged male *Mecp2-null* mice show a significant reduction in the expression of (G) Population C marker *Cartpt*; and a significant increase in the expression of Population B marker *Nfia* when compared to tissues of age-and sex-matched littermate wildtype (WT) control mice (** p < 0.01, **** p < 0.0001; Students’ t-test). By contrast, the expression of Nfix shows no statistically significant differences between the two genotypes (Students’ t-test).

### Loss of BDNF – TrkB signaling results in a loss of *Vip* expression but not in expression of VIP receptor genes or *Cartpt* in the adult murine small intestinal myenteric plexus

VIPergic interneuron dysfunction has been thought to drive cortical dysfunction in *Mecp2-null* mice and thus, in RTT patients. Given MeCP2’s role as a transcriptional regulator, it can be argued that MeCP2 loss directly affects *Vip* expression in the ENS. Alternately, it can be argued that congenital reduction in BDNF levels and the resulting reduction in TrkB signaling contributes to a significant reduction in *Vip* expression in the small intestinal ENS. We hypothesized that loss of *Vip* expression in *Mecp2-null* mice is a result of congenital reduction in BDNF – TrkB signaling. To test this, we used the *Wnt1*-cre:*TrkB^fl/fl^* adult male mice, where the TrkB.FL isoform is conditionally and congenitally knocked out in all neural crest-derived cells in a *Wnt1*-cre dependent manner. We have previously shown that these mice show a selective loss of TrkB.FL (but not of TrkB.T1) isoform in the small intestinal myenteric plexus to cause a loss of enteric BDNF – TrkB signaling and a significant reduction in whole GI motility without alterations to the expression of *Uchl1, Nos1*, or *Chat* genes in their LM-MP tissues [15]. Here, we performed qRT-PCR using primers specific to *Vip, Vipr1,* and *Vipr2* and found that similar to the *Mecp2-null* mice, the small intestinal LM-MP tissues of male *Wnt1*-cre:*TrkB^fl/fl^* also showed a significant reduction in the expression of *Vip*, but no change in the expression of the VIP receptor genes (n = 4/group, mean ± S.E. fold change of *Vip*: WT: 1.006 ± 0.064; *Wnt1*-cre:*TrkB^fl/fl^*: 0.719 ± 0.014, p = 0.004; mean ± S.E. fold change of *Vipr1*: WT: 1.079 ± 0.266; *Wnt1*-cre:*TrkB^fl/fl^*: 1.692 ± 0.365, p = 0.22; mean ± S.E. fold change of *Vipr2*: WT: 1.011 ± 0.086; *Wnt1*-cre:*TrkB^fl/fl^*: 1.853 ± 1.033, p = 0.44; Student’s t-test; **Fig 8 A-C).**

**Figure 8:**
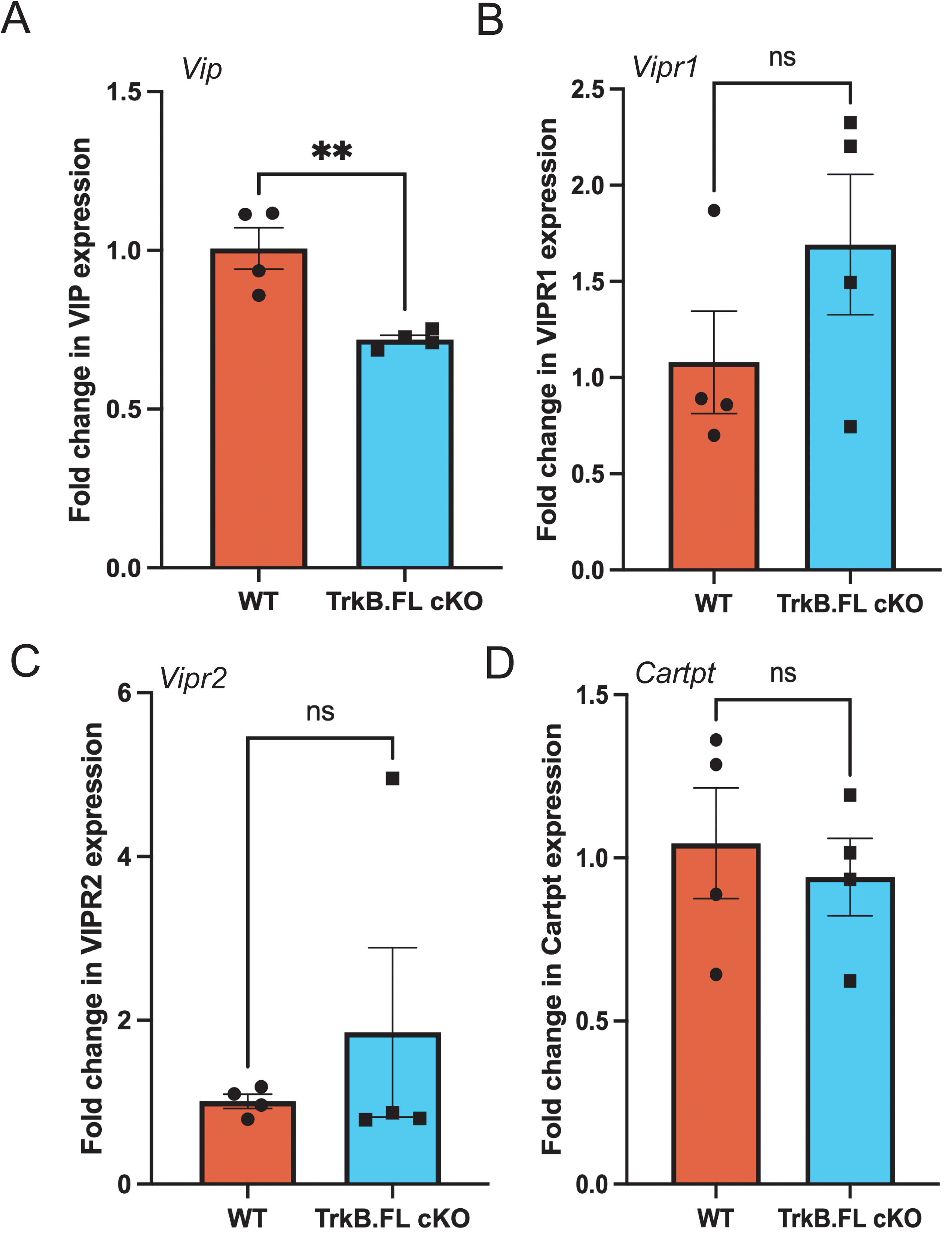
Loss of TrkB signaling in small intestinal neural crest-derived cells does not completely recapitulate. MeCP2-loss associated changes to VIPergic signaling genes qRT-PCR – based analyses of relative abundance of expressed transcripts from small intestinal longitudinal muscle – myenteric plexus (LM-MP) tissues of *Wnt1*-cre:*TrkB^fl/fl^* mice, where TrkB.FL is conditionally knocked out from neural crest-derived enteric neurons (TrkB.FL cKO), and age-matched sex-matched littermate control wildtype (WT) mice show that (A) expression of inhibitory neuropeptide *Vip* is significantly reduced in the TrkB.FL cKO tissues compared to WT tissues (** p < 0.01; Students’ t-test). However, neither the expression of (B) *Vipr1*, nor the expression of (C) *Vipr2* is significantly different between the two genotypes. (D) Further, the expression of *Cartpt,* which is expressed by the *Nos1^+^ Vip1^+^ Cartpt^+^* Subpopulation C of the inhibitory neuronal cluster is not found to be significantly different between the two genotypes (** p < 0.01, ns p > 0.05; Students’ t-test).

We tested whether loss of TrkB.FL from neural crest cells in ENS causes a significant reduction in expression of *Cartpt* as observed in the *Mecp2-null* murine tissues. However, unlike the *Mecp2-null* mice, adult male *Wnt1*-cre:*TrkB^fl/fl^* mice showed no statistically significant change in *Cartpt* gene expression (n = 4/group, mean ± S.E. fold change of *Cartpt*: WT: 1.04 ± 0.16; *Wnt1*-cre:*TrkB^fl/fl^*: 0.94 ± 0.12, p = 0.63; Student’s t-test; **Fig 8D).**

### Pharmacological modulation of TrkB regulates *Vip* expression in wildtype and *Mecp2*-null ENS cells

Since reduced *Bdnf* transcript expression was detectable at an earlier age than *Vip* downregulation, we tested whether pharmacological TrkB inhibition reduced *Vip* expression in differentiating postnatal ENS cultures. We and others have previously tested and shown that differentiating cultures of post-natal ENS cells can be used to assess neuronal differentiation [30–32]. This would allow us to further test whether TrkB signaling regulates *Vip* in the ENS. To that, we tested whether pharmacological inhibition of TrkB (to mimic reduced *Bdnf* levels) significantly altered *Nos1* and *Vip* expression. We found that differentiating post-natal murine ENS cells exposed to ANA-12, a specific antagonist of TrkB [33–35], showed significant downregulation of *Vip* and *Cartpt* but not of *Nos1*, suggesting again that suppressing TrkB signaling does not alter the expression of the larger inhibitory (nitrergic) marker but specifically targets *Vip^+^ Cartpt^+^* neuronal population (n = 4/group, mean ± S.E. fold change of *Nos1*: Control: 1.027 ± 0.136; ANA-12: 0.886 ± 0.092, p = 0.42; mean ± S.E. fold change of *Vip*: Control: 1.001 ± 0.022; ANA-12: 0.761 ± 0.036, p = 0.001; mean ± S.E. fold change of *Cartpt*: Control: 1.001 ± 0.026; ANA-12: 0.706 ± 0.054, p = 0.002; Student’s t-test; Student’s t-test; **Fig 9 A, B**).

**Figure 9:**
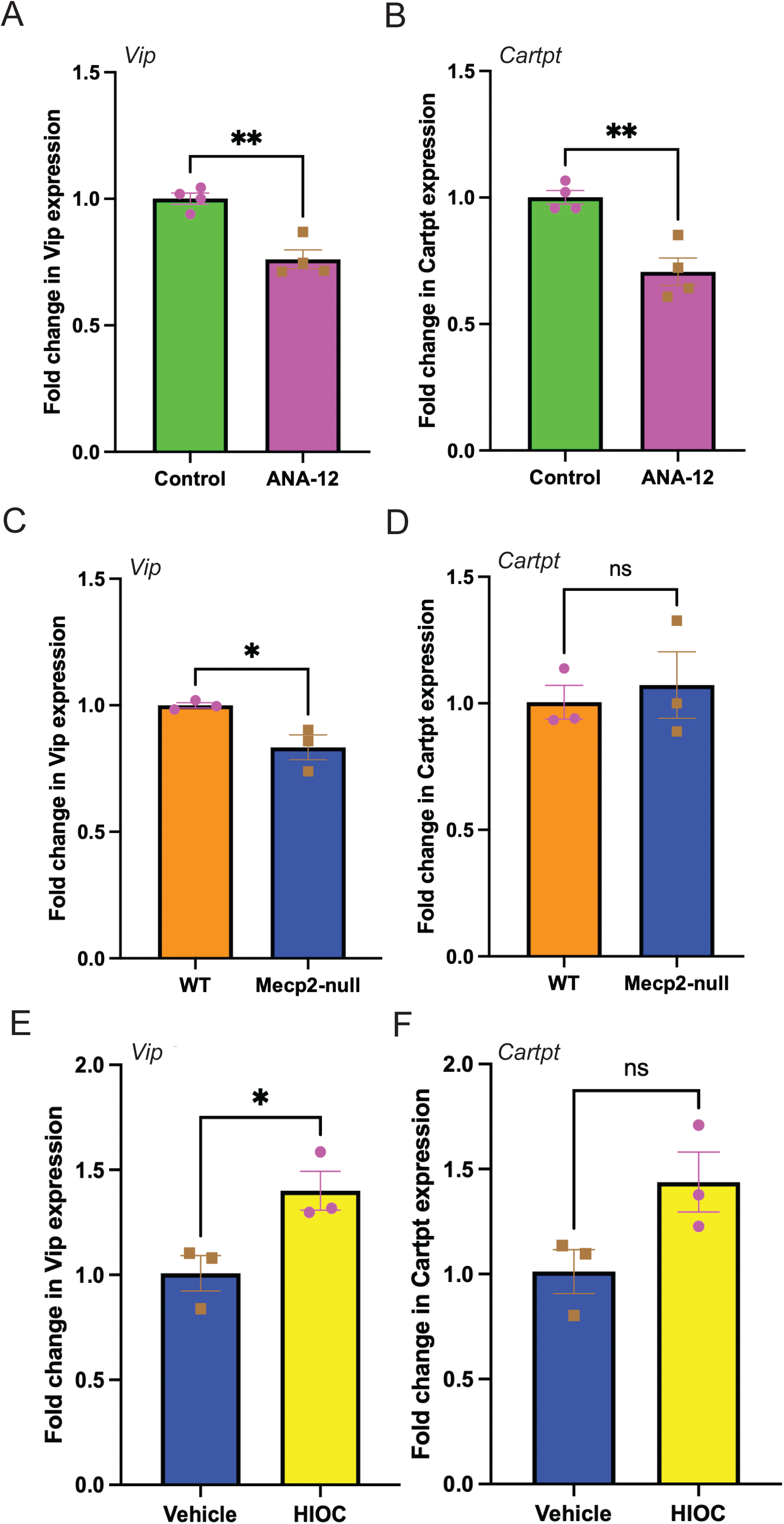
Pharmacological modulation of TrkB regulates Vip expression in wildtype and Mecp2-null ENS cells. *In vitro* differentiating post-natal murine small intestinal ENS cell cultures from wildtype mice when cultured with and without exposure to TrkB antagonist ANA-12 shows that ANA-12 exposure over a course of 14 days leads to a significant decline in the expression of (A) *Vip* and (B) *Cartpt*, when compared to ENS cells differentiated without ANA-12. Differentiating post-natal murine small intestinal ENS cell cultures from wildtype (WT) mice and *Mecp2-null* mice show that *Mecp2* loss drives a significant decline in the expression of (C) *Vip* but no change in expression of (D) *Cartpt*, when compared to ENS cells derived from WT mice. *In vitro* differentiating post-natal murine small intestinal ENS cell cultures from *Mecp2-null* mice when cultured with and without exposure to TrkB agonist HIOC shows that HIOC exposure over a course of 7 days leads to a significant increase in the expression of (E) *Vip* and a non-significant increase in expression of (F) *Cartpt*, when compared to cells differentiated without HIOC. (*p < 0.05, ** p < 0.01; Student’s t-test).

We next tested whether differentiating ENS cells from P35 *Mecp2*-null mice also differed in the expression of *Vip* and *Cartpt* when compared to littermate control mice. *Mecp2-*null mice ENS cell cultures showed progressive cell loss in culture beyond day 7, and thus, for these experiments, we performed a curtailed differentiation protocol of 7 days, instead of the 14 days used for prior experiments. In this experiment, we again found no difference in expression of *Nos1*, but a significant decrease in expression of *Vip*. *Cartpt* expression, however, shows no statistically significant difference after 7 days of differentiation (n = 3/group, mean ± S.E. fold change of *Nos1*: WT: 1.008 ± 0.868; *Mecp2-null*: 0.900 ± 0.077, p = 0.40; mean ± S.E. fold change of *Vip*: WT: 1.000 ± 0.001; *Mecp2-null*: 0.833 ± 0.048, p = 0.029; mean ± S.E. fold change of *Cartpt*: WT: 1.004 ± 0.066; *Mecp2-null*: 1.072 ± 0.131, p = 0.66; Student’s t-test; **Fig 9C, D**).

Given that *Mecp2-null* ENS cells recreate the reduced expression of *Vip in vitro*, we next tested whether pharmacological stimulation of TrkB in the differentiating Mecp2-null cells promotes the expression of *Vip* and *Cartpt*, which would suggest that TrkB signaling can promote the genesis of *Vip* and *Cartpt*-expressing neuronal population. *Mecp2-*null mice ENS cell cultures when differentiated in presence of HIOC, a TrkB agonist we have used previously [15] showed significant increase in expression of *Vip* and a strong tendency towards increased expression of *Cartpt* when compared to cells that were cultured with vehicle (Control), suggesting that continued stimulation of TrkB during neuronal differentiation promotes the genesis of the *Vip^+^* inhibitory neuronal population in 7 days of differentiation, despite the loss of *Mecp2* (n = 3/group, mean ± S.E. fold change of *Vip*: Control: 1.008 ± 0.084; HIOC: 1.400 ± 0.092, p = 0.035; mean ± S.E. fold change of *Cartpt*: Control: 1.012 ± 0.104; HIOC: 1.438 ± 0.142, p = 0.07; Student’s t-test; **Fig 9 E, F**).

## Discussion

Although RTT patients suffer debilitating GI dysmotility, a lack of mechanistic understanding of these disorders has led to a lack of therapies that can improve overall GI function in these patients. In this study, we identify reduced enteric *Bdnf* expression in *Mecp2*-null male mice before the emergence of detectable GI dysmotility. At P55, delayed transit accompanies reduced *Vip* and *Cartpt* expression and increased *Nfia* expression, without detectable changes in overall neuronal density or nitrergic neuronal abundance. These findings suggest selective alterations in inhibitory neuronal phenotypes or population composition rather than generalized neuronal depletion. Conditional deletion of TrkB.FL in neural crest-derived cells reproduces the reduction in *Vip*, while pharmacological TrkB inhibition reduces *Vip* and *Cartpt* expression in differentiating ENS cultures. While ENS cultured cells from *Mecp2-null* mice show significant reduction of Vip, thus recreating the *in vivo* phenotype associated with GI dysmotility, TrkB stimulation of these cultures increases *Vip* expression. Together, these findings implicate reduced enteric BDNF-TrkB signaling as a mediator of dysfunctional VIPergic phenotype that is associated with evolving GI dysmotility in a MeCP2 loss model of RTT. The results support the identification of TrkB and VIP as potential points of therapeutic intervention for GI dysmotility in RTT patients with loss of MeCP2 function.

MeCP2 is an X-linked gene and most of the RTT patients are females, as males with MeCP2 mutations die early after birth [36]. However, since male *Mecp2* mutant hemizygous mice that have only a single copy of X-linked *Mecp2* gene, effectively making them *Mecp2-null*, show a faster regression than female *Mecp2* mutant hemizygous mice [37–39], we used male mice for this study to map out the effects of MeCP2 loss on ENS and GI motility.

Contrary to prior report that showed an increase in expression of NOS1 protein in the ENS of *Mecp2-null* mice [7], P55-aged male *Mecp2-null* mice that suffered from significant GI dysmotility did not show any significant alteration in either the numbers of Hu-immunolabeled neurons in the small intestinal myenteric plexus, nor in the numbers of the inhibitory NOS1-immunolabeled nitrergic neurons. The absence of significant changes in *Nos1* expression, together with comparable NOS1-immunolabeled neuronal densities, argues against a broad alteration in nitrergic neuronal abundance at P55.

MeCP2 is known to regulate *Bdnf* transcription from exon 4, and hence *Bdnf IV* isoform [11, 40, 41]. Our data show lower expression of *Bdnf IV,* and *VI* transcripts in small intestinal LM-MP from P55 *Mecp2-null* males. Since *Bdnf* transcripts *IV* and *VI* account for approximately 90% of enteric *Bdnf* expression [15], their reduction suggests a substantial decrease in the dominant transcript pool. Neither TrkB.FL nor TrkB.T1 transcript expression differed significantly between genotypes at P35 or P55, suggesting limited ligand availability in the intestines of the *Mecp2-null* mice. Our previous work implicates enteric BDNF–TrkB signaling in glucocorticoid-associated dysmotility [15]; the present findings raise the possibility that related mechanisms contribute to the phenotype in *Mecp2-null* mouse model of RTT.

However, compared to our earlier work, where a decrease in BDNF in adult mice led to an acute slowdown of GI motility, in the *Mecp2-null* mice, *Bdnf IV* and *VI* expression was already reduced at P35 age where WGTT was not significantly altered. This suggests that the relationship between BDNF availability and WGTT in juvenile mice is distinct from our observation at the adult stage. By P55, reduced *Vip* expression accompanied delayed whole gut transit. These cross-sectional comparisons place show that reduced *Bdnf* transcript expression precedes significant *Vip* downregulation and delayed transit.

In addition to alterations in BDNF expression, the *Mecp2-null* RTT model also shows a dysregulation in VIPergic system in the CNS [21, 42]. Given that VIP is an important neuropeptide that regulates intestinal motility, epithelial cell biology, and gut immunity and microbiota [27, 28, 43], we tested and found that *Vip* expression is significantly downregulated and the expression of its receptors *Vipr1* and *Vipr2* is significantly upregulated in the ENS of P55-aged *Mecp2-null* male mice. *Vipr1* (also called VPAC1R) is broadly expressed by intestinal cells, including enteric neurons, immune cells, [44–48] and *Vipr2* (also known as VPAC2R), which drives relaxation of gut musculature, is predominantly found in intestinal smooth muscle cells [49, 50]. Thus, the resulting aberrant VIPergic signaling in the ENS of *Mecp2-null* male mice has implications beyond altered GI motility, as it may also help explain the microbial dysbiosis observed in RTT mice models and patients [51–55]. Furthermore, *Cartpt* encoded neuropeptide protein Cocaine and Amphetamine Transcript; CART is abundantly present in the ENS and plays a role in maintaining tissue architecture and in regulation of blood glucose [56, 57]. Loss of *Cartpt* expression in ENS of *Mecp2-null* mice thus provides a possible mechanism through which metabolic complications of RTT patients can be explained [58].

Reduced *Bdnf* expression preceding delayed whole gut transit time and reduced enteric *Vip* expression suggested a link between TrkB signaling and *Vip* expression in the ENS. Using a mouse model *Wnt1*-cre:*TrkB^fl/fl^*, where the full-length isoform of TrkB (which carries out BDNF signaling) is knocked out in all neural crest-derived cells, including those of the ENS, conditional TrkB.FL deletion was associated with lower enteric *Vip* expression, without significant differences in VIP receptor transcripts. We have previously used this mouse model to show that such a congenital loss of TrkB in neural crest-derived neurons of the ENS does not cause any significant change to total neuronal numbers or to proportions of nitrergic neurons [15]. Here, we use this mouse model to show that reduced BDNF – TrkB signaling drives reduced expression of *Vip* in the adult murine small intestinal myenteric plexus. The culture-based *in vitro* experiments extend this evidence by demonstrating that enteric *Vip* expression remains pharmacologically responsive to TrkB modulation. ANA-12 reduced *Vip* and *Cartpt* expression without significantly altering *Nos1*, supporting selective regulation of these neuropeptide-associated programs rather than uniform suppression of the nitrergic phenotype. Unlike congenital TrkB.FL deletion, ANA-12 also reduced *Cartpt*, suggesting that developmental context, treatment duration, or culture conditions influence the response. It is important to note that while the congenital deletion acts only on TrkB.FL, while leaving TrkB.T1 levels unchanged in the ENS [15], ANA-12 binds the extracellular pocket of TrkB, which is both conserved between TrkB.FL and TrkB.T1 [59]. Thus, the lack of effect on *Cartpt* in the conditional TrkB.FL deletion mouse model and the effect of pan-TrkB blockage on *Cartpt in vitro* suggests that TrkB.T1’s role may not be restricted to be a BDNF sink, and that it may be involved in a distinct signaling system in the ENS. Prior reports from the Tessarollo lab provide evidence of such signaling in other organs [60], the parallels between those studies and TrkB signaling in the ENS needs to be further investigated.

Within a model of ongoing postnatal and adult enteric neurogenesis [16, 17], our data on TrkB signaling-driven VIP signal suggests that TrkB may help differentiating precursors acquire and maintain specific neuronal identities, especially the VIP^+^ CARTPT^+^ phenotype. Interpreted alongside the inhibitory neuronal subtypes resolved in the Southard-Smith datasets, our findings suggest that reduced TrkB activity could selectively impair the replenishment or maintenance of VIPergic neurons, progressively compromising their representation despite preserved overall nitrergic neuronal abundance. This mechanism could explain why reduced *Bdnf* expression precedes detectable *Vip* downregulation and delayed transit. Conversely, the increase in *Vip* expression following HIOC treatment suggests that continued postnatal neuronal plasticity may provide a therapeutic opportunity to restore deficient neuronal phenotypes through TrkB stimulation. Establishing whether this response reflects neuronal differentiation, survival, or renewed neuropeptide expression will require lineage tracing and cellular quantification.

Although our results suggest a loss of a VIP and CART-expressing neuronal population in *Mecp2-null* mice and a downregulation of *Vip* expression but not loss of a specific neuronal subpopulation in *Wnt1*-cre:*TrkB^fl/fl^*male mice, establishing these require further dedicated experiments. Simple immunolabeling of VIP and CART in the small intestinal myenteric plexus tissue will not provide sufficient clarity to distinguish between the possibility of loss of VIP expression in neurons compared to loss of subclass of neurons that express VIP. Thus, future studies will focus on utilizing fate-mapping mice where VIPergic neurons are indelibly labeled with reporters to study whether loss of MeCP2 drives loss of VIPergic cells or of expression of *Vip* from these cells. In culture, lineage-resolved measurements of differentiation, neuronal abundance, and viability will be particularly important because *Mecp2-null* preparations developed progressive cell loss beyond day 7, necessitating a shorter differentiation protocol than that used for ANA-12 experiments. This difference limits us from testing how a longer stimulation with HIOC may increase the development of CART-expressing and other neuronal subpopulations in *Mecp2-null* cells.

Most individuals with RTT are females heterozygous for a pathogenic *MECP2* variant, whereas our experiments used hemizygous *Mecp2-null* males. The onset and progression of GI dysfunction, and the conservation of the BDNF–TrkB and VIPergic abnormalities identified here, therefore remain to be established in heterozygous females. Their slower disease course may also provide a longer window for testing whether sustained TrkB stimulation preserves enteric neuronal phenotypes and improves GI function.

Our findings support a model in which early enteric BDNF deficiency contributes to later VIPergic dysfunction and GI dysmotility following MeCP2 loss. The responsiveness of *Mecp2-null* ENS cultures to TrkB stimulation identifies a therapeutic avenue for further investigation, with restoration of neuronal function and GI motility remaining key tests of its potential.

## Acknowledgements

This work was supported by funding from NIA R01AG066768, R21AG072107, and Pilot grant from the Harvard Digestive Disease Core (SK). This work was also supported by funding from the TUBITAK 2214-A International Research Fellowship Program (GS), Maryland Genetics, Epidemiology, and Medicine training program sponsored by the Burroughs Welcome Fund (JS), NIGMS grant T32GM148383 (JS), and by Walter Benjamin Fellowship (528835020) from Deutsche Forschungsgemeinschaft (PS).

## Supplementary Figure

**Figure S1:** Schematic of TrkB isoform-specific primers. Schematic shows the location of the primers that allow us to target TrkB.FL isoform (which contains exons 11 and 13 but not exon 12) and TrkB.T1 isoform (which contains exons 11 and 12 but none thereafter) differentially.

## References

1. Liyanage, V.R. and M. Rastegar, Rett syndrome and MeCP2. Neuromolecular Med, 2014. 16(2): p. 231–64.

2. Neul, J.L., et al., Rett syndrome: revised diagnostic criteria and nomenclature. Ann Neurol, 2010. 68(6): p. 944–50.

3. Ip, J.P.K., N. Mellios, and M. Sur, Rett syndrome: insights into genetic, molecular and circuit mechanisms. Nature Reviews Neuroscience, 2018. 19(6): p. 368–382.

4. Motil, K.J., et al., Gastrointestinal and nutritional problems occur frequently throughout life in girls and women with Rett syndrome. J Pediatr Gastroenterol Nutr, 2012. 55(3): p. 292–8.

5. Sharkey, K.A. and G.M. Mawe, The enteric nervous system. Physiological Reviews, 2023. 103(2): p. 1487–1564.

6. Rao, M. and M.D. Gershon, The bowel and beyond: the enteric nervous system in neurological disorders. Nat Rev Gastroenterol Hepatol, 2016. 13(9): p. 517–28.

7. Wahba, G., et al., Activity and MeCP2-dependent regulation of nNOS levels in enteric neurons. Neurogastroenterol Motil, 2016. 28(11): p. 1723–1730.

8. Wahba, G., et al., MeCP2 in the enteric nervous system. Neurogastroenterol Motil, 2015. 27(8): p. 1156–61.

9. Zhou, Z., et al., Brain-specific phosphorylation of MeCP2 regulates activity-dependent Bdnf transcription, dendritic growth, and spine maturation. Neuron, 2006. 52(2): p. 255–69.

10. Sampathkumar, C., et al., Loss of MeCP2 disrupts cell autonomous and autocrine BDNF signaling in mouse glutamatergic neurons. eLife, 2016. 5: p. e19374.

11. Chang, Q., et al., The disease progression of Mecp2 mutant mice is affected by the level of BDNF expression. Neuron, 2006. 49(3): p. 341–8.

12. Boesmans, W., et al., Brain-derived neurotrophic factor amplifies neurotransmitter responses and promotes synaptic communication in the enteric nervous system. Gut, 2008. 57(3): p. 314–22.

13. Bonfiglio, F., et al., GWAS of stool frequency provides insights into gastrointestinal motility and irritable bowel syndrome. Cell Genom, 2021. 1(3): p. None.

14. Chen, F., et al., Brain-derived neurotrophic factor accelerates gut motility in slow-transit constipation. Acta Physiol (Oxf), 2014. 212(3): p. 226–38.

15. Slosberg, J., and Puttapaka S.N. et al., Stress-induced glucocorticoid signaling impairs enteric neurotrophin BDNF-TrkB pathway and drives gastrointestinal dysmotility. J Biol Chem. 302(7):113095.

16. Gorecki, A.M., et al., *Detection of mitotic neuroblasts provides additional evidence of steady state neurogenesis in the adult small intestinal myenteric plexus*. eneuro, 2025: p. ENEURO.0005-24.2025.

17. Kulkarni, S., et al., Adult enteric nervous system in health is maintained by a dynamic balance between neuronal apoptosis and neurogenesis. Proc Natl Acad Sci U S A, 2017. 114(18): p. E3709–E3718.

18. Benthal, J.T., A.A. May-Zhang, and E.M. Southard-Smith, *Building consensus: construction of a juvenile and adult scRNA-seq meta-atlas for dataset* comparisons *and harmonizing transcriptomic definitions of enteric neuron subtypes*. BMC Genomics, 2026. 27(1): p. 50.

19. Barde, Y.A., The physiopathology of brain-derived neurotrophic factor. Physiol Rev, 2025. 105(4): p. 2073–2140.

20. Honore, S.M., et al., Neuronal loss and abnormal BMP/Smad signaling in the myenteric plexus of diabetic rats. Auton Neurosci, 2011. 164(1-2): p. 51–61.

21. Li, Q., et al., Circadian rhythm disruption in a mouse model of Rett syndrome circadian disruption in RTT. Neurobiology of Disease, 2015. 77: p. 155–164.

22. Hamnett, R., et al., Regional cytoarchitecture of the adult and developing mouse enteric nervous system. Current Biology, 2022. 32(20): p. 4483–4492.e5.

23. Lelievre, V., et al., Gastrointestinal dysfunction in mice with a targeted mutation in the gene encoding vasoactive intestinal polypeptide: a model for the study of intestinal ileus and Hirschsprung’s disease. Peptides, 2007. 28(9): p. 1688–99.

24. Shi, X.Z. and S.K. Sarna, Gene therapy of Cav1.2 channel with VIP and VIP receptor agonists and antagonists: a novel approach to designing promotility and antimotility agents. Am J Physiol Gastrointest Liver Physiol, 2008. 295(1): p. G187–g196.

25. Shi, X.Z. and S.K. Sarna, Homeostatic and therapeutic roles of VIP in smooth muscle function: myo-neuroimmune interactions. Am J Physiol Gastrointest Liver Physiol, 2009. 297(4): p. G716–25.

26. Bai, X., et al., Vasoactive Intestinal Polypeptide Plays a Key Role in the Microbial-Neuroimmune Control of Intestinal Motility. Cell Mol Gastroenterol Hepatol, 2024. 17(3): p. 383–398.

27. Fung, C., et al., VPAC1 receptors regulate intestinal secretion and muscle contractility by activating cholinergic neurons in guinea pig jejunum. American Journal of Physiology-Gastrointestinal and Liver Physiology, 2014. 306(9): p. G748–G758.

28. Jakob, M.O., et al., Enteric nervous system-derived VIP restrains differentiation of LGR5+ stem cells toward the secretory lineage impeding type 2 immune programs. Nature Immunology, 2025. 26(12): p. 2227–2243.

29. May-Zhang, A.A., et al., Combinatorial Transcriptional Profiling of Mouse and Human Enteric Neurons Identifies Shared and Disparate Subtypes In Situ. Gastroenterology, 2021. 160(3): p. 755–770.e26.

30. Becker, L., et al., Divergent fate and origin of neurosphere-like bodies from different layers of the gut. Am J Physiol Gastrointest Liver Physiol, 2012. 302(9): p. G958–65.

31. Silva, A.T., et al., Neural progenitors from isolated postnatal rat myenteric ganglia: expansion as neurospheres and differentiation in vitro. Brain Res, 2008. 1218: p. 47–53.

32. Guyer, R.A., et al., Single-cell multiome sequencing clarifies enteric glial diversity and identifies an intraganglionic population poised for neurogenesis. Cell Rep, 2023. 42(3): p. 112194.

33. Zhang, H. and Z. Cai, Inhibition of miR-206 Ameliorates Seizures and Cognitive Deficits via BDNF/TrkB Signaling in Pilocarpine-Induced TLE Mice. Brain Sci, 2026. 16(10).

34. Sims, S.K., et al., Intranasal Inhibition of Tropomyosin Receptor Kinase B Signaling during Early Postnatal Development Disrupts Growth and Sensorimotor Maturation in Neonatal Rodents. Dev Neurosci, 2026: p. 1–11.

35. Uju, C. and S. Unniappan, Brain derived neurotrophic factor promotes oocyte maturation through the TrkB-MAPK-PI3K pathway in zebrafish. Mol Cell Endocrinol, 2026. 612: p. 112703.

36. Ribeiro, M.C. and J.L. MacDonald, Sex differences in Mecp2-mutant Rett syndrome model mice and the impact of cellular mosaicism in phenotype development. Brain Research, 2020. 1729: p. 146644.

37. Samaco, R.C., et al., Female Mecp2(+/-) mice display robust behavioral deficits on two different genetic backgrounds providing a framework for pre-clinical studies. Hum Mol Genet, 2013. 22(1): p. 96–109.

38. Lang, M., et al., Rescue of behavioral and EEG deficits in male and female Mecp2-deficient mice by delayed Mecp2 gene reactivation. Hum Mol Genet, 2014. 23(2): p. 303–18.

39. Lombardi, L.M., S.A. Baker, and H.Y. Zoghbi, MECP2 disorders: from the clinic to mice and back. J Clin Invest, 2015. 125(8): p. 2914–23.

40. Li, W. and L. Pozzo-Miller, BDNF deregulation in Rett syndrome. Neuropharmacology, 2014. 76 **Pt C**(0 0): p. 737–46.

41. Sun, Y.E. and H. Wu, The Ups and Downs of BDNF in Rett Syndrome. Neuron, 2006. 49(3): p. 321–323.

42. Mossner, J.M., et al., Developmental loss of MeCP2 from VIP interneurons impairs cortical function and behavior. eLife, 2020. 9: p. e55639.

43. Pirzgalska, R.M., et al., Neuroepithelial VIP–VIPR1 interactions differentially control enteric type 1 and type 2 immunity. Nature Immunology, 2025. 26(12): p. 2244–2255.

44. Jayawardena, D., et al., Expression and localization of VPAC1, the major receptor of vasoactive intestinal peptide along the length of the intestine. American Journal of Physiology-Gastrointestinal and Liver Physiology, 2017. 313(1): p. G16–G25.

45. Yukawa, T., et al., Differential expression of vasoactive intestinal peptide receptor 1 expression in inflammatory bowel disease. Int J Mol Med, 2007. 20(2): p. 161–7.

46. Cocco, E., et al., The Expression of Vasoactive Intestinal Peptide Receptor 1 Is Negatively Modulated by MicroRNA 525-5p. PLOS ONE, 2010. 5(8): p. e12067.

47. Sun, W., et al., Altered expression of vasoactive intestinal peptide receptors in T lymphocytes and aberrant Th1 immunity in multiple sclerosis. International Immunology, 2006. 18(12): p. 1691–1700.

48. Fung, C., et al., VPAC1 receptors regulate intestinal secretion and muscle contractility by activating cholinergic neurons in guinea pig jejunum. Am J Physiol Gastrointest Liver Physiol, 2014. 306(9): p. G748–58.

49. Harmar, A.J., et al., Distribution of the VPAC2 receptor in peripheral tissues of the mouse. Endocrinology, 2004. 145(3): p. 1203–10.

50. Mahavadi, S., et al., Caveolae-dependent internalization and homologous desensitization of VIP/PACAP receptor, VPAC, in gastrointestinal smooth muscle. Peptides, 2013. 43: p. 137–45.

51. Strati, F., et al., Altered gut microbiota in Rett syndrome. Microbiome, 2016. 4(1): p. 41.

52. Neier, K., et al., Sex disparate gut microbiome and metabolome perturbations precede disease progression in a mouse model of Rett syndrome. Communications Biology, 2021. 4(1): p. 1408.

53. Talbot, J., et al., Feeding-dependent VIP neuron-ILC3 circuit regulates the intestinal barrier. Nature, 2020. 579(7800): p. 575–580.

54. Al Nabhani, Z., V. Thevin, and L. Morelli, Neuronal VIP wires the intestinal epithelial cell function. Mucosal Immunology, 2026. 19(1): p. 1679–1681.

55. Jacobson, A., et al., The intestinal neuro-immune axis: crosstalk between neurons, immune cells, and microbes. Mucosal Immunology, 2021. 14(3): p. 555–565.

56. Ekblad, E., CART in the enteric nervous system. Peptides, 2006. 27(8): p. 2024–2030.

57. Muller, P.A., et al., Microbiota-modulated CART(+) enteric neurons autonomously regulate blood glucose. Science, 2020. 370(6514): p. 314–321.

58. Kyle, S.M., N. Vashi, and M.J. Justice, Rett syndrome: a neurological disorder with metabolic components. Open Biol, 2018. 8(2).

59. Cazorla, M., et al., Identification of a low–molecular weight TrkB antagonist with anxiolytic and antidepressant activity in mice. The Journal of Clinical Investigation, 2011. 121(5): p. 1846–1857.

60. Tessarollo, L. and S. Yanpallewar, TrkB Truncated Isoform Receptors as Transducers and Determinants of BDNF Functions. Front Neurosci, 2022. 16: p. 847572.

