## Supplementary Figure 1 for "Reduced enteric *Bdnf* expression and TrkB-dependent regulation of *Vip* in a *Mecp2-null* mouse model of Rett syndrome-associated gastrointestinal dysmotility"

### Mouse Ntrk2 (TrkB) isoform transcript structure and primer locations

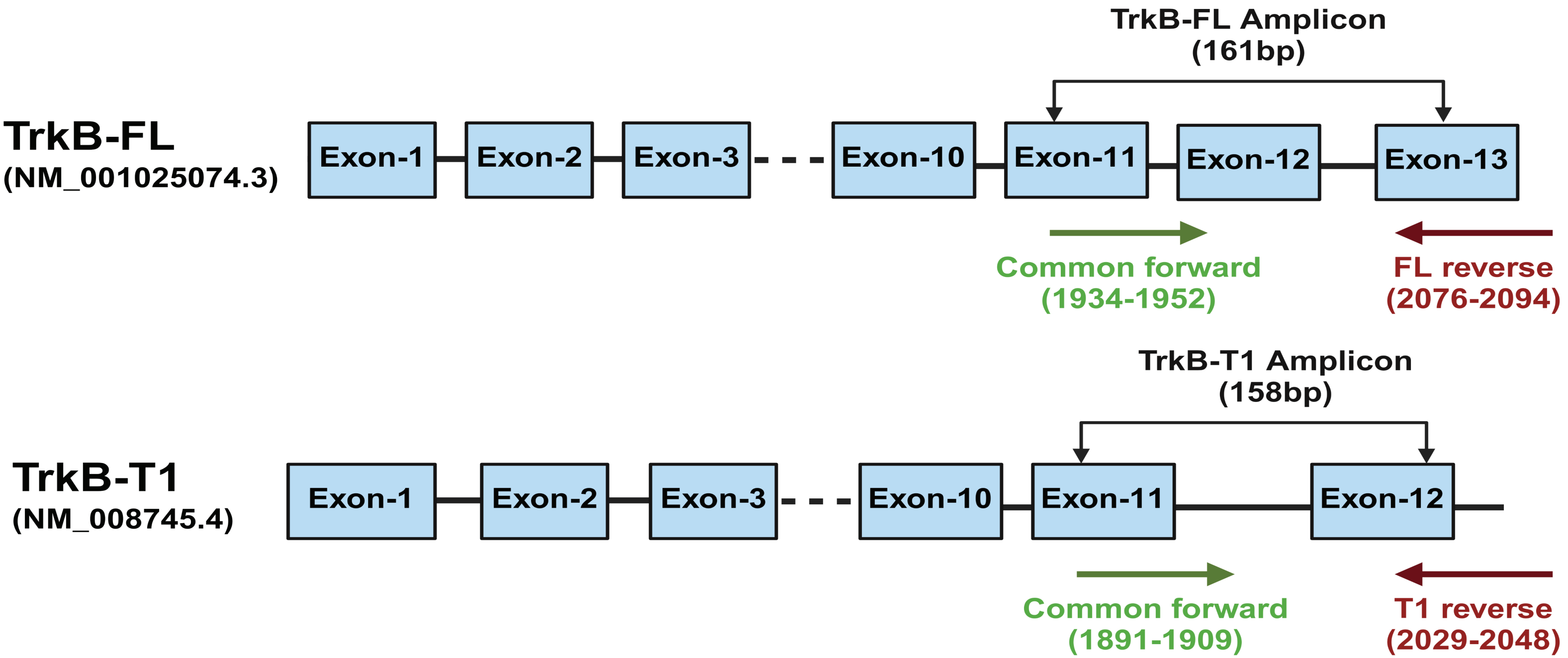


Coordinates are relative to the NM_001025074.3 (TrkB-FL) and NM_008745.4 (TrkB-T1) transcripts.
